# Single-cell transcriptomics reveal diverging pathobiology and opportunities for precision targeting in scleroderma-associated versus idiopathic pulmonary arterial hypertension

**DOI:** 10.1101/2024.10.25.620225

**Authors:** Tijana Tuhy, Julie C. Coursen, Tammy Graves, Michael Patatanian, Christopher Cherry, Shannon E. Niedermeyer, Sarah L. Khan, Darin T. Rosen, Michael P. Croglio, Mohab Elnashar, Todd M. Kolb, Stephen C. Mathai, Rachel L. Damico, Paul M. Hassoun, Larissa A. Shimoda, Karthik Suresh, Micheala A. Aldred, Catherine E. Simpson

**Affiliations:** Division of Pulmonary and Critical Care Medicine, Johns Hopkins University, Baltimore, MD, USA; Division of Pulmonary Medicine, Indiana University, Indianapolis, IN, USA; Faculty of Medicine, Ain Shams University, Cairo, Egypt; Division of Pulmonary and Critical Care Medicine, University of Miami, Miami, FL, USA

## Abstract

**Introduction:** Pulmonary arterial hypertension (PAH) involves progressive cellular and molecular change within the pulmonary vasculature, leading to increased vascular resistance. Current therapies targeting nitric oxide (NO), endothelin, and prostacyclin pathways yield variable treatment responses. Patients with systemic sclerosis-associated PAH (SSc-PAH) often experience worse outcomes than those with idiopathic PAH (IPAH).

**Methods:** Lung tissue samples from four SSc-PAH, four IPAH, and four failed donor specimens were obtained from the Pulmonary Hypertension Breakthrough Initiative (PHBI) lung tissue bank. Single-cell RNA sequencing (scRNAseq) was performed using the 10X Genomics Chromium Flex platform. Data normalization, clustering, and differential expression analysis were conducted using Seurat. Additional analyses included gene set enrichment analysis (GSEA), transcription factor activity analysis, and ligand-receptor signaling. Pharmacotranscriptomic screening was performed using the Connectivity Map.

**Results:** SSc-PAH samples showed a higher proportion of fibroblasts and dendritic cells/macrophages compared to IPAH and donor samples. GSEA revealed enriched pathways related to epithelial-to-mesenchymal transition (EMT), apoptosis, and vascular remodeling in SSc-PAH samples. There was pronounced differential gene expression across diverse pulmonary vascular cell types and in various epithelial cell types in both IPAH and SSc-PAH, with epithelial to endothelial cell signaling observed. Macrophage to endothelial cell signaling was particularly pronounced in SSc-PAH. Pharmacotranscriptomic screening identified TIE2, GSK-3, and PKC inhibitors, among other compounds, as potential drug candidates for reversing SSc-PAH gene expression signatures.

**Discussion:** Overlapping and distinct gene expression patterns exist in SSc-PAH versus IPAH, with significant molecular differences suggesting unique pathogenic mechanisms in SSc-PAH. These findings highlight the potential for precision-targeted therapies to improve SSc-PAH patient outcomes. Future studies should validate these targets clinically and explore their therapeutic efficacy.

---

Pulmonary arterial hypertension (PAH) is characterized by progressive cellular and molecular changes within the pulmonary vasculature leading to structural remodeling and rising pulmonary vascular resistance.^1,2^ Until recently, approved therapies have targeted one of three molecular pathways: the nitric oxide (NO) signaling pathway, the endothelin pathway, or the prostacyclin pathway.^3,4^ With these therapies, treatment responses have been modest and variable, with patients experiencing a heterogeneity of treatment effects.^5^ In particular, patients with PAH related to systemic sclerosis (SSc-PAH) derive less benefit and experience worse clinical outcomes compared to other PAH subgroups.^6–8^

We and others have shown that in SSc-PAH, pathophysiologic differences at the molecular, cellular, and tissue levels underlie observed differences in clinical outcomes.^9–14^ A detailed understanding of the molecular differences between SSc-PAH and idiopathic PAH (IPAH) might allow for exploitation of aberrant signaling pathways unique to SSc-PAH (and/or present in both of these PAH subtypes) for drug targeting. Recently, sotatercept, a novel ligand-trap molecule that targets TGF-beta signaling in PAH, improved hemodynamics and walk distance for patients already treated with drugs targeting NO, endothelin, and/or prostacyclin signaling.^15,16^ The success of sotatercept demonstrates that targeting unaddressed molecular aberrancies can yield additional clinical benefit.

Most previous omics investigations in PAH have examined circulating plasma or serum.^17–21^ With this study, we sought to identify distinct and overlapping gene expression patterns in SSc-PAH versus IPAH human lung tissue. To maximize insights, we took a single-cell approach to RNA sequencing, in contrast to the bulk sequencing performed in most previous PAH studies.^12,22,23^ We leveraged cell-specific gene expression patterns to identify small molecule compounds with high potential to reverse disease-specific signatures, as a means of prioritizing candidates for drug repositioning.

## Study Design and Methods

Complete methods are in the Online Supplement. All data and materials have been made publicly available at the Gene Expression Omnibus and can be accessed at [https://www.ncbi.nlm.nih.gov/gds/persistent – specific URL pending acceptance]. Tissue collection and analysis was approved by Institutional Review Boards at all sites contributing to PHBI. The current study makes use of de-identified samples thus does not constitute human subjects’ research.

## Results

From the Pulmonary Hypertension Breakthrough Initiative (PHBI) lung tissue bank, we identified four SSc-PAH, four IPAH, and four failed donor specimens judged suitable to serve as controls (Table 1). Features detected, UMIs, and percentage mitochondrial DNA per sample are shown in Supplemental Figures 1a and 1b. After quality control, 13,650 cells were analyzed. After batch correction (Supplemental Figure 2), clustering on integrated samples resulted in 16 transcriptionally distinct groups of cells, which were manually annotated as shown in Figure 1a. Top DEGs by cluster are shown in the heatmap in Figure 1b. Cell types were hierarchically clustered as shown in Figure 1c. When manually annotated clusters were projected onto LungMap, the cluster manually labeled “proliferative” aligned with LungMap’s deuterosomal cell cluster. The cluster labeled “hematopoietic” aligned with megakaryocytes/platelets. Other manually annotated clusters aligned with similarly annotated LungMap clusters.

**Table 1.** Demographic information and sample identifiers for subjects donating lung tissue to the Pulmonary Hypertension Breakthrough Initiative (PHBI) tissue bank.

| Sample | Age | Sex | Race | Diagnosis | mPAP<br>(mmHg) | PVR<br>(WU) | CI<br>(L/min/m<sup>2</sup>) | FVC<br>(% pred) | DLCO<br>(% pred) |
| --- | --- | --- | --- | --- | --- | --- | --- | --- | --- |
| BA-044 | 30YR | Male | White | FDL |  |  |  |  |  |
| BA-048 | 17YR | Male | White | FDL |  |  |  |  |  |
| BA-053 | 50YR | Female | Unknown | FDL |  |  |  |  |  |
| BA-058 | 54YR | Male | Unknown | FDL |  |  |  |  |  |
| BA-024 | 54YR | Female | White | SSc-PAH | 55 | 8.23 | 3.36 | 93 | 38 |
| BA-025 | 64YR | Female | Unknown | SSc-PAH | 32 | 5.98 | 3.13 | 85 | 25 |
| ST-005 | 79YR | Female | White | SSc-PAH | 42 | 8.12 | 2.26 | 81 | 35 |
| ST-059 | 66YR | Female | White | SSc-PAH | 57 | 12.29 | 4.07† | 94 | 28 |
| BA-008 | 7YR | Male | White | IPAH | 60 | ND | 3.41 | 72 | ND |
| BA-014 | 41YR | Female | Unknown | IPAH | 55 | 13.06 | 2.2 | 66* | 61 |
| CC-030 | 63YR | Female | White | IPAH | 58 | 10 | 3.1 | 86 | 73 |
| ST-042 | 53YR | Male | White | IPAH | 45 | 8.6 | 2.16 | 113 | 97 |

Definition of abbreviations. mPAP: mean pulmonary arterial pressure; PVR: pulmonary vascular resistance; CI: cardiac index; FVC:
forced vital capacity; DLCO: diffusing capacity of the lung for carbon monoxide; FDL: failed donor lung; SSc-PAH: systemic sclerosis-associated pulmonary arterial hypertension; IPAH: idiopathic pulmonary arterial hypertension; ND: values not collected
\* Contemporaneously measured TLC was 82% predicted.
† Reflects cardiac output. Cardiac index not collected.
All PAH patients who contributed tissue samples were exposed to combination therapy with PDE5 inhibitors (PDE5-i), endothelin receptor antagonists (ERA), and prostacyclin agents (PCA) leading up to transplant (with exception of ST-005, who received PDE5-i, PCA, and a third investigational drug at the time of explant).

**Figure 1.**
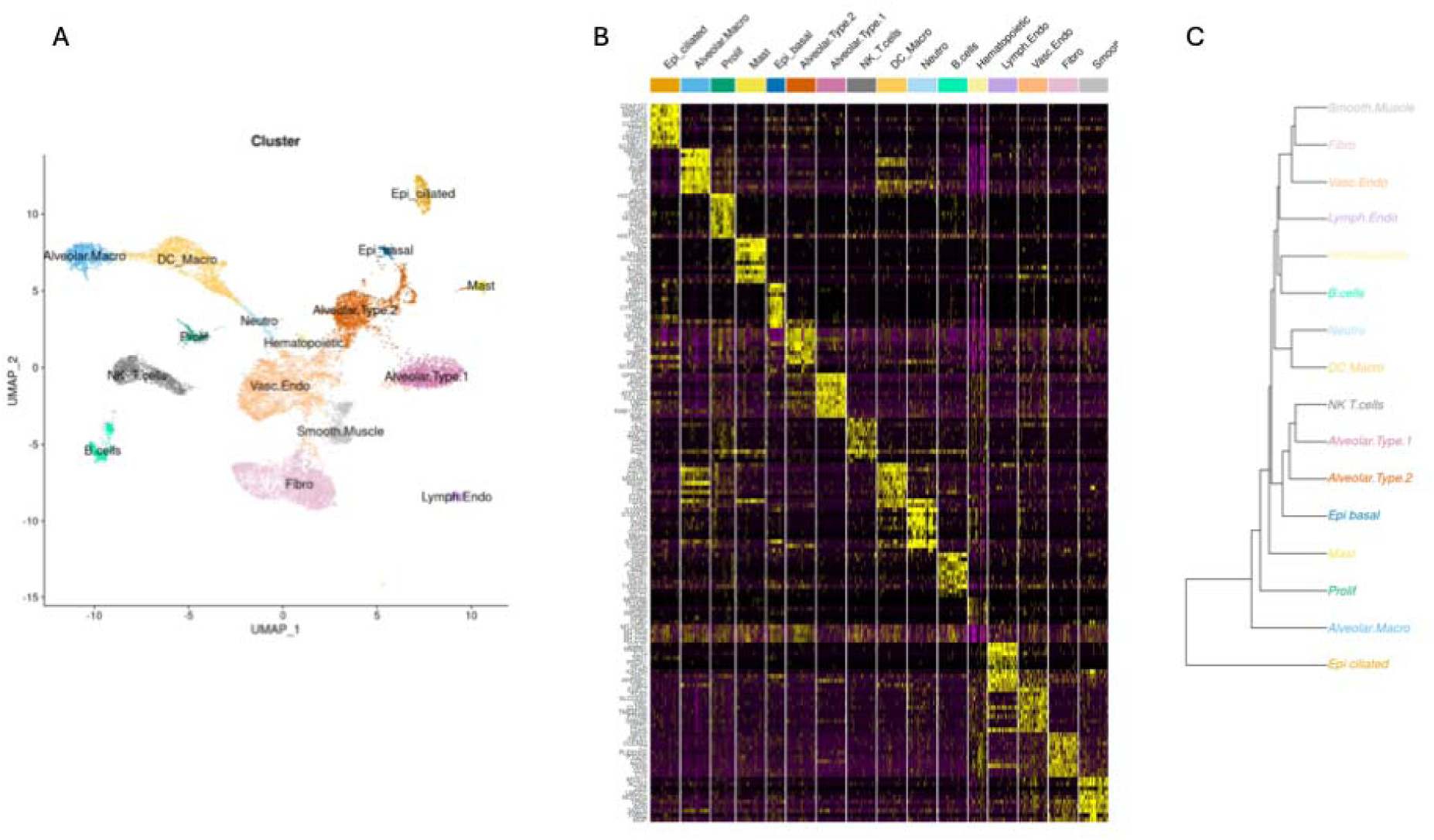
Overview of clustering results. A) Uniform manifold approximation and projection (UMAP) of human lung scRNAseq annotated and colored by cell cluster. B) Purple-yellow heatmap of top-most differentially expressed genes by cluster. Cells for a given cluster are compared to all other cells in the dataset. Yellow cells represent upregulated genes, while purple cells represent downregulated genes. C) Hierarchical dendrogram of clustered cell types.

### Cluster proportions

Comparisons of cell type proportions across conditions (e.g., SSc-PAH, IPAH, or donor) are shown in Supplemental Figures 3a and 3b. SSc-PAH fibroblasts contributed a large proportion of cells to the SSc-PAH total cell count, representing approximately 20% of SSc-PAH cells, compared to approximately 10% fibroblasts present in IPAH and donor cell populations. SSc-PAH dendritic cells/macrophages represented a higher proportion of SSc-PAH cells compared to IPAH cells. IPAH and SSc-PAH smooth muscle cell (SMC) proportions were numerically higher compared to donors (with borderline p-values 0.08 and 0.17, respectively). By contrast, SSc-PAH vascular endothelial cells contributed a lower proportion of the total SSc-PAH cell count (approximately 10% versus 20% in IPAH and donors, with borderline p-value 0.099).

### Differential expression and GSEA

A topologic overview of differentially expressed genes (DEGs) by cell cluster is depicted for each PAH subtype (relative to donors) in Figure 2a. Figure 2b shows DEG counts by cluster. Overall, DEGs in SSc-PAH SMCs and fibroblasts showed greater fold-change differences compared to IPAH cells. Across cell types, there tended to be a higher number of DEGs from SSc-PAH samples compared to IPAH samples. For both SSc-PAH and IPAH, vascular endothelial cells demonstrated the highest number of DEGs.

**Figure 2.**
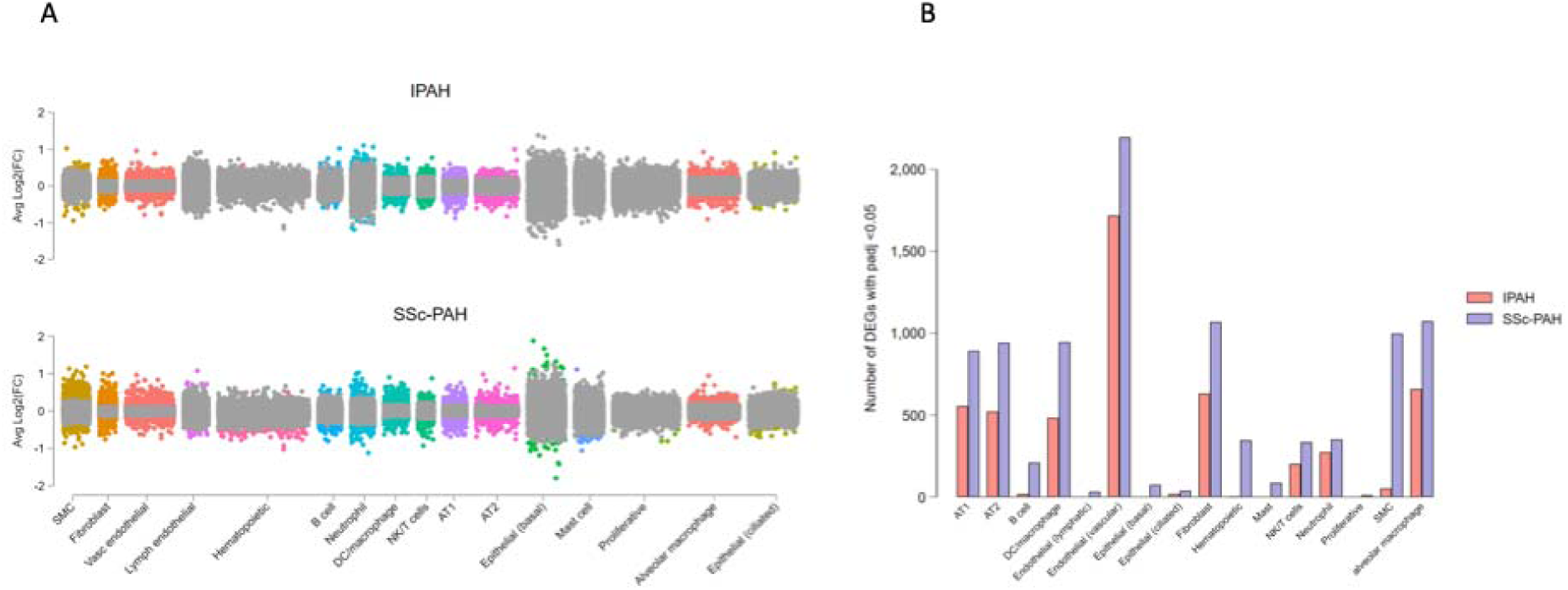
Differential expression analysis. A) Jitter plot showing differentially expressed genes for each cell type in IPAH vs controls (top) and SSc-PAH vs controls (bottom). Each dot represents the average log2FC of a gene. Dots indicating differentially expressed genes that surpass a false discovery corrected p-value <0.05 are colored by cell type. Gray dots indicate genes that did not surpass the threshold for statistical significance. B) Bar plot representing counts of differentially expressed genes (relative to donors) for IPAH and SSc-PAH cell clusters.

Genes demonstrating highly significant differential expression (e.g., adjusted *p-*value <1×10^-4^) are shown for each PAH subtype vs. donors, and for SSc-PAH vs. IPAH, in Supplemental Figures 3-11. Several genes associated with PAH were differentially expressed across diverse cell types, such as *IGFBP7*^29,30^ (upregulated across vascular endothelial cells, fibroblasts, SMCs, AT1 (alveolar type 1) and AT2 (alveolar type 2) cells); *COL18A1*^31–33^ (upregulated in vascular endothelial cells and fibroblasts); and *SERPINE1*^34^ (upregulated in vascular endothelial cells, fibroblasts, SMCs, AT2 cells).

In vascular endothelial cells, genes encoding IGF binding proteins (IGFBPs), *ID3*, and *DEPP1* were among the most differentially expressed genes in both SSc-PAH and IPAH; IGFBPs were more upregulated in SSc-PAH relative to IPAH. In both PAH subtypes, fibroblasts demonstrated upregulation of genes encoding various structural proteins, including collagens, laminin, and MMPs, and downregulation of ADAMTS proteins. SMC expression patterns were more distinct between disease subtypes, with IPAH SMCs demonstrating differential expression of *CCN2, ID3,* and *NOTCH3*, and SSc-PAH SMCs demonstrating greater differential expression of genes involved in extracellular matrix organization, cell adhesion, and tissue remodeling, including *FN1, IGFBP7, VIM*, and *LTBP2*.

In both SSc-PAH and IPAH, AT1 and AT2 cells demonstrated differential expression of genes associated with PAH. Genes involved in extracellular matrix organization (e.g., *LMNA, FN1*) and vascular remodeling (e.g., *TNFSF12, VEGFA*) were differentially expressed in AT1 cells. Genes involved in lipid metabolism (*FASN, LPCAT1*) and tissue remodeling (*SERPINA3*) were differentially expressed in AT2 cells. In both AT1 and AT2 cells, genes encoding surfactant proteins A, B, and C were upregulated in IPAH and SSc-PAH, with differential gene expression more pronounced in SSc-PAH. In airway epithelial cells, differential expression of genes encoding mucins, including *MUC5AC* and *MUC5B*, was observed.

GSEA results are presented for IPAH vs. donor cells in Fig 3a, and for SSc-PAH vs. donor cells in Fig 3b. In IPAH vs. donor cells, significant enrichments were observed in pathways governing inflammation and inflammatory responses, including TNF-alpha signaling, interleukin signaling, and interferon responses. Gene sets regulated by the oncogenes KRAS and Myc, which influence cellular proliferation and apoptosis, were enriched. Pathways involved in vascular remodeling were also enriched, including Mtorc1 signaling and TGF-beta signaling. Many of the same enrichments were observed in GSEA of SSc-PAH vs. donors (Fig 3b). In SSc-PAH, gene sets involved in epithelial to mesenchymal transition (EMT) were enriched across multiple cell types, most significantly in vascular endothelial cells and SMCs, but also in fibroblasts, epithelial cells, and AT2 cells. Gene sets implicated in myogenesis were uniquely enriched in SSc-PAH endothelial cells, SMCs, and fibroblasts. SSc-PAH SMCs, epithelial cells, and alveolar macrophages demonstrated enrichments across multiple gene sets (Fig 3b).

**Figure 3.**
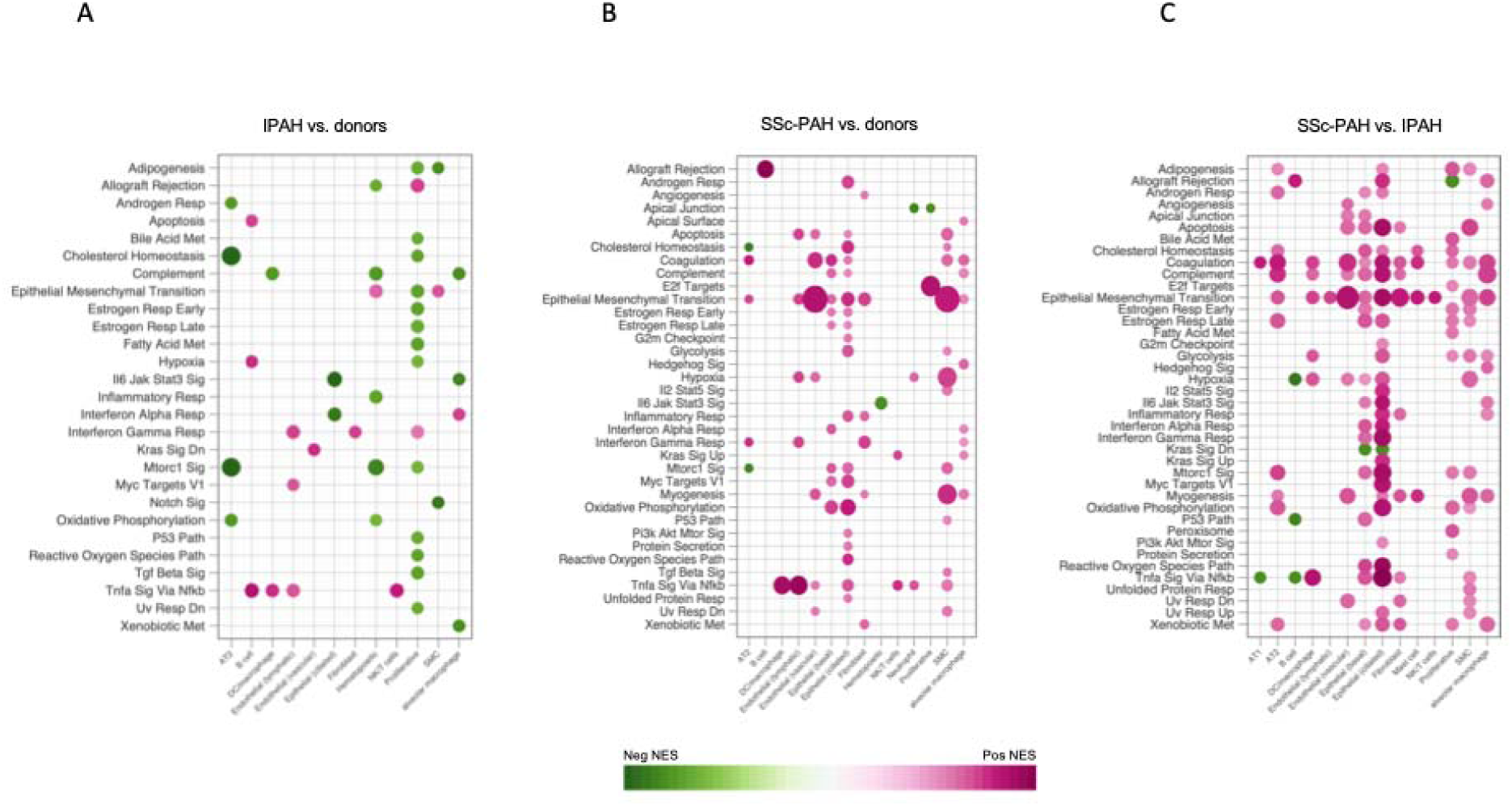
Gene set enrichment analysis. Heatmaps showing cluster-specific gene set enrichment analysis of gene signatures from IPAH (A) and SSc-PAH (B) samples compared with donor controls using hallmark pathways from the Molecular Signatures Database. SSc-PAH vs IPAH GSEA is shown in (C). Gene sets are listed on the on the y-axis, and cell clusters are on the x-axis. The dot size corresponds to −log10(p-value), and color represents the normalized enrichment score (NES) from GSEA, indicating upregulation (pink) or downregulation (purple). NES is provided for enriched gene sets with FDR-adjusted p-value <0.05.

When we performed head-to-head GSEA of SSc-PAH vs. IPAH samples, gene expression patterns associated with EMT were significantly enriched in SSc-PAH across multiple cell types, including vascular endothelial cells, lymphatic endothelial cells, SMCs, fibroblasts, AT2 cells, and epithelial cells (Fig 3c). A similar pattern of more pronounced enrichment across SSc-PAH cell types was observed for apoptosis, cholesterol homeostasis, coagulation, complement, and myogenesis, among others. Additionally, multiple gene sets were enriched in SSc-PAH airway and alveolar epithelial cells relative to IPAH.

### Overlapping and distinct expression patterns in SSc-PAH vs. IPAH

As shown in Supplemental Figure 12, there was little overlapping differential gene expression for B cells, neutrophils, smooth muscle cells, and ciliated epithelial cells, however cell totals were lower for these clusters than for other cell types. There was more substantial overlap in DEGs for vascular endothelial cells, fibroblasts, natural killer (NK)/T cells, and macrophages.

To assess for consistency in the directionality and magnitude of differential gene expression common to both PAH subtypes, we took the intersect of common DEGs and plotted log2FC differences in IPAH vs. SSc-PAH (Supplemental Fig 13a and b). For several clusters, including SMCs, fibroblasts, and epithelial cells, the position of the line of best fit relative to equality suggested that, on average, the magnitude of differential gene expression was greater for SSc-PAH than for IPAH. For vascular endothelial cells, the magnitude of average differential expression was similar in IPAH and SSc-PAH.

### Gene regulatory analysis and intercellular signaling

To infer gene regulatory differences between PAH subgroups, we performed differential TF activity analysis for each cell cluster using UCell (Supplemental Fig 14). Clusters with the most significant differential TF activity included SMCs, fibroblasts, vascular endothelial cells, and AT1 cells. The top 25 TFs with differential activity for each cell type are displayed in Figure 4. Among IPAH cells, HOXB7, a TF implicated in promoting angiogenesis and regulating genes involved in cell proliferation and survival, was among the most significantly upregulated TFs, with increased activity in vascular endothelial cells (Figure 4). In SSc-PAH samples, EGFR and RARB, both of which regulate genes involved in cell differentiation and survival, were among the most significantly upregulated TFs. EGFR signaling has been previously implicated in PAH.^35,36^ Other TFs implicated in PAH^37^ were among top hits, including PPARGC1A,^38,39^ HMGA1,^40^ NFKB,^41^ and TFAM.^42^ Though not among the top 25 for any cluster, several other TFs implicated in PAH were differentially active in PAH cells, including SMAD3, KLF4, and HIF-1a.^37^

**Figure 4.**
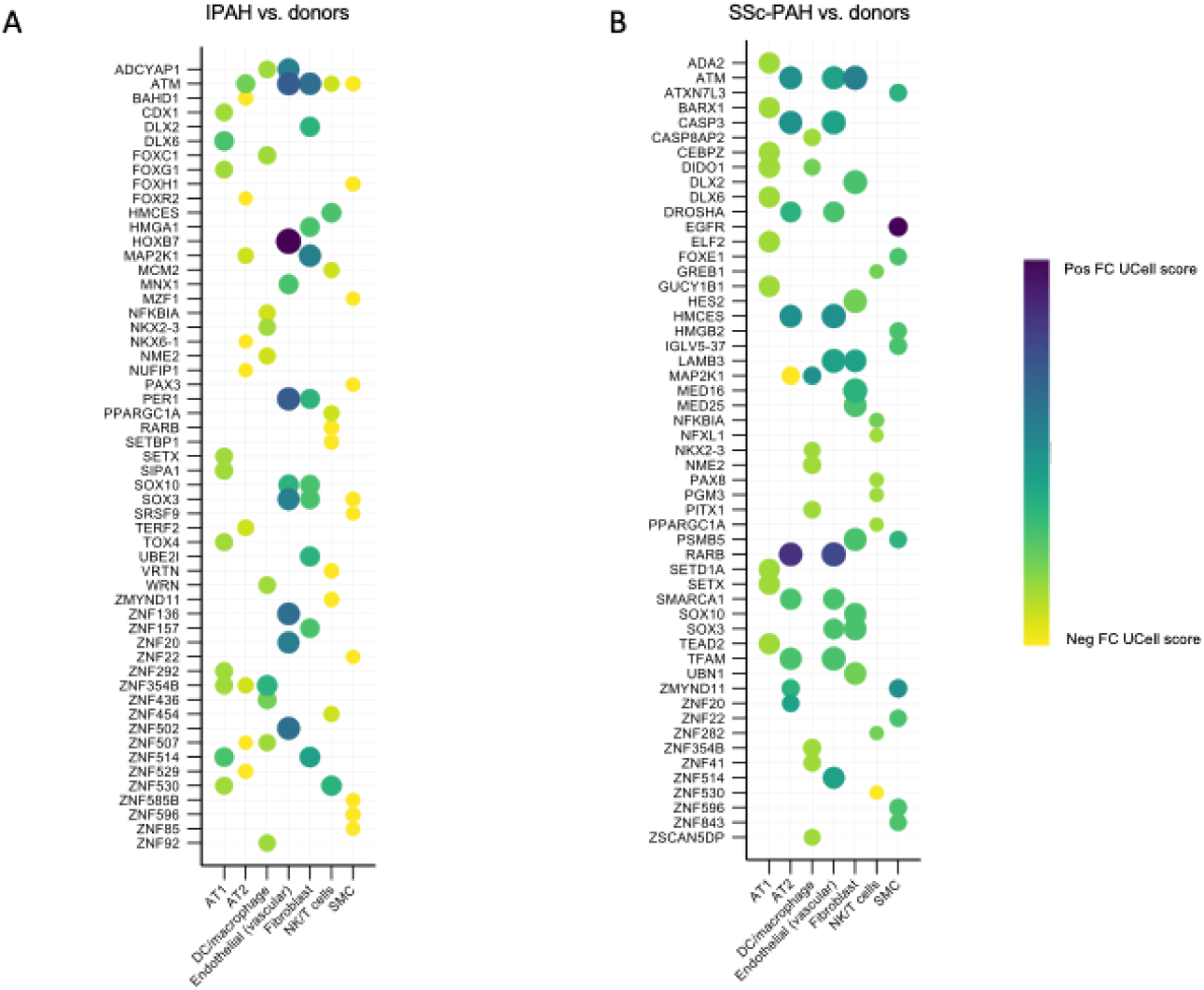
Gene regulatory analysis of differential transcription factor activity. Heatmaps showing the fold-change differences in UCell scores by condition for the top 25 transcription factors (sorted by significance) that are most differentially active for each cluster for IPAH vs donors (A) and SSc-PAH vs donors (B). Dot color represents the FC difference between conditions, with purple indicating upregulation and yellow indicating downregulation of TF activity relative to donors. Dot size corresponds to −log10(p-value) of the FC difference.

We combined our assessment of differential TF activity with scored ligand-receptor interactions to infer a global understanding of intercellular signaling. Signaling patterns for each PAH subtype are depicted in the chord diagrams in Fig 5A and Fig 5B, respectively. In general, SSc-PAH intercellular patterns were characterized by increased signaling (relative to IPAH) to vascular endothelial cell receptors from diverse cell types, including alveolar macrophages, AT1 cells, and smooth muscle cells. The heatmap in Figure 5C quantifies inferred signaling differences between SSc-PAH and IPAH. The most pronounced differences were observed in SMC and alveolar macrophage signaling to vascular endothelial cells (greater in SSc-PAH) and in AT1 and alveolar macrophage signaling to hematopoietic cells (greater in IPAH).

**Figure 5.**
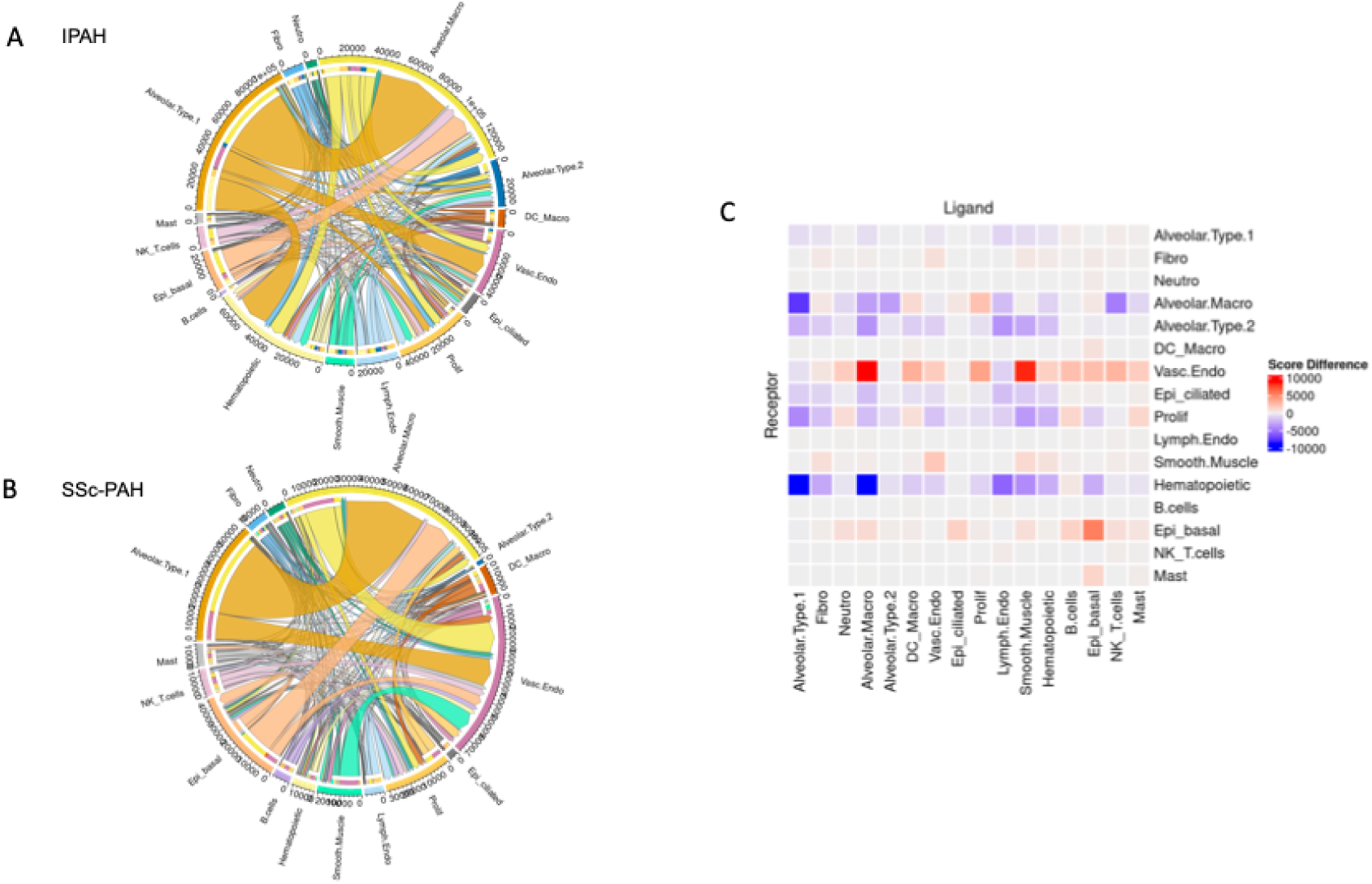
Global intercellular signaling patterns. Chord diagrams showing intercellular inferences via CellPhoneDB for IPAH (A) and SSc-PAH (B) cells. Each segment around the circle corresponds to a different cell cluster identified in the dataset. The length of colored segments indicates the relative abundance of each cell type. Chords connecting the segments represent ligand-receptor interactions, with the thickness of each chord proportional to the interaction strength. Colors are used to distinguish between different cell types and their interactions. Arrowheads indicate directionality from ligand to receptor. C) Heatplot of quantitative differences in ligand-receptor interaction scores in SSc-PAH versus IPAH.

In both SSc-PAH and IPAH, common signaling interactions were inferred between AT1 cells and basal epithelial cells and alveolar macrophages, and between these same epithelial cell types and vascular endothelial cells. Signaling from AT1 cells to vascular endothelial cells was characterized most prominently by VEGFA-FLT1 ligand-receptor binding in SSc-PAH, and by TGFB2-TGFB2 receptor2 ligand-receptor binding in IPAH (Supplemental Figure 15). Inferred interactions between AT1 cells and alveolar macrophages consisted of CD47 – SIRB1 complex ligand-receptor signaling, with activation of various downstream transcription factors, including MAPK3 and NR1D1. Interactions between basal epithelial cells and alveolar macrophages consisted predominantly of IL33-IL33 receptor ligand-receptor signaling.

In SSc-PAH, SMC signaling to vascular endothelial cells consisted of PGF-FLT1 complex ligand-receptor binding and TGF-beta signaling (Supplemental Figure 16). Alveolar macrophage signaling to vascular endothelial cells consisted of VEGFB-FLT1 complex and TGFB1-TGF beta receptors 1 and 2 (Supplemental Figure 17). Chord diagrams depicting ligand-receptor interactions coupled to downstream TF activity for various cell types are shown in supplemental Figure 18. Violin plots depicting expression of commonly implicated signaling molecules by condition are shown in supplemental Figures 19A and 19B.

### Pharmacotranscriptomic screening

To infer potential druggable targets, and to investigate the potential for precision targeting of SSc-PAH, we input cluster-specific gene expression signatures into CMap to identify pharmacologic compounds with opposing (e.g., negative) gene expression signatures. Dot plots summarizing connectivity scores for top-most opposing compounds in IPAH and SSc-PAH, across all cell lines tested in CMAP, are shown in Supplemental Figure 20A. Top candidates for reversing SSc-PAH signatures included BRD-A92800748, a TIE2 inhibitor; BRD-K04923131, a GSK-3 inhibitor that influences JAK/STAT signaling; THZ-2-98-01, an inhibitor of IRAK1, which induces upregulation of the TF NFKB; GW-5074, a selective c-Raf inhibitor; and enzastaurin, a PKC-beta inhibitor known to reverse experimental PH in the Sugen hypoxia rodent model.^43^ Average connectivity scores for tested compounds, grouped by mechanism of action, are shown for IPAH and SSc-PAH in Supplemental Figures 20B and 20C respectively. When analyzed by mechanism of action, TIE inhibitors, GSK inhibitors, and PKC inhibitors showed highest potential for reversing SSc-PAH gene signatures.

To focus our assessment, because abnormal endothelial cell biology is a cardinal feature of PAH,^1,44^ we filtered our initial results for compounds with opposing signatures in human umbilical vein endothelial cell lines (HUVECs) in particular. Figure 6A and Table 2 provide an overview of the top compounds tested in HUVECs against SSc-PAH. The TIE2 inhibitor BRD-A92800748 gave the most negative connectivity score, with c-score −0.93 for the fibroblast cluster, and c-score −0.86 for the vascular endothelial cell cluster. Other compounds with strongly opposing connectivity scores in HUVECs against SSc-PAH included TGF-beta receptor inhibitors, PKC inhibitors, and bromodomain inhibitors (Table 2).

**Figure 6.**
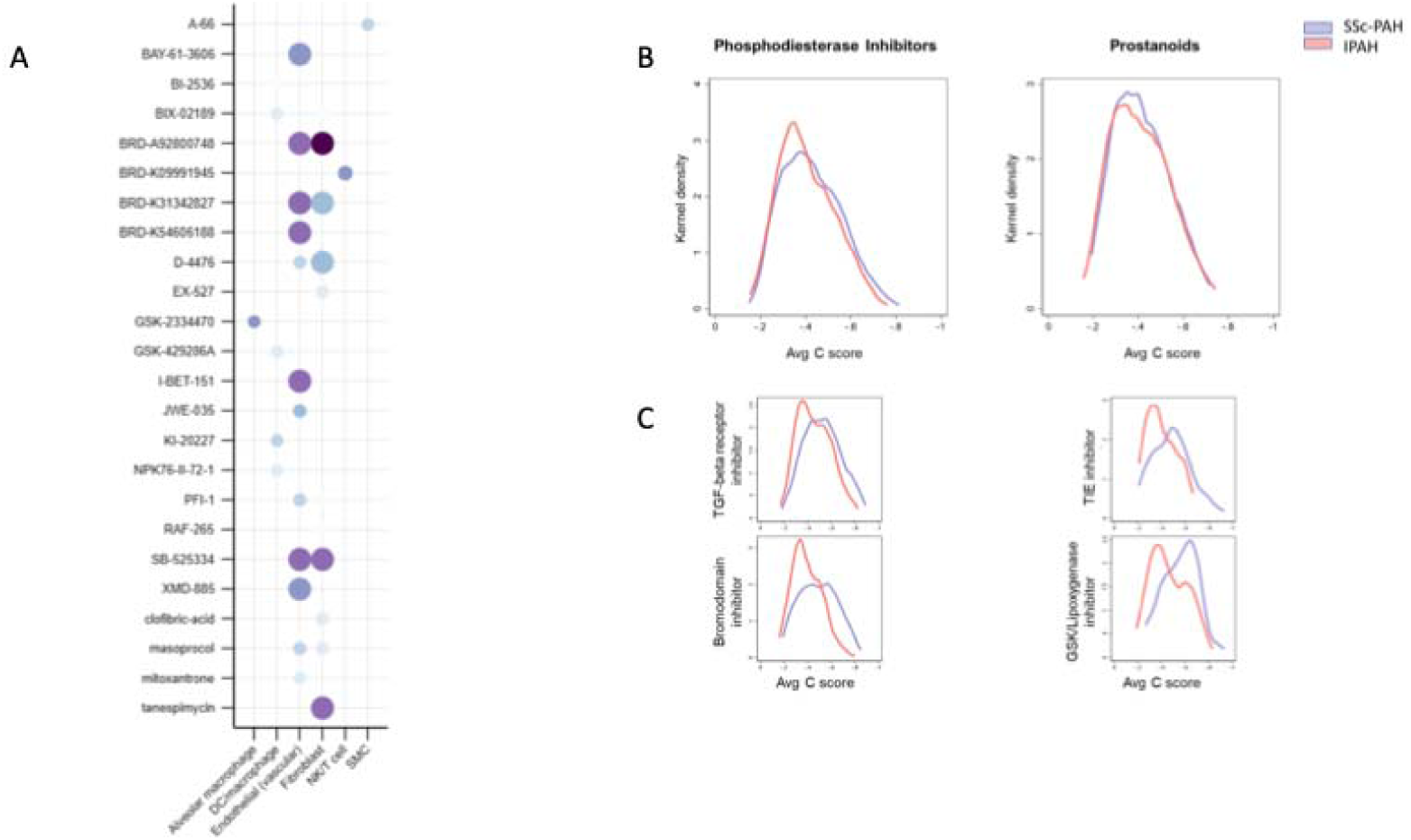
Summary of pharmacotranscriptomic screen. A) Heatmap of connectivity scores for the compounds opposing cluster-specific gene expression signatures from SSc-PAH cells when tested in human umbilical vein endothelial cell lines. Individual compound names are on the y-axis, and cell types are on the x-axis. Scores are depicted for compounds with FDR-corrected p-value <0.05. Dot color corresponds to connectivity score, with dark purple representing −1, and light blue representing −0.7. Dot size corresponds to −log10(p-value). B) Kernel density plots depicting the distribution of connectivity scores opposing SSc-PAH gene expression signatures (in purple) and IPAH gene expression signatures (in pink) for approved and commonly used pulmonary hypertension therapies. B)) Kernel density plots depicting the distribution of connectivity scores opposing SSc-PAH gene expression signatures (in purple) and IPAH gene expression signatures (in pink) for investigational pulmonary hypertension therapies.

**Table 2.**
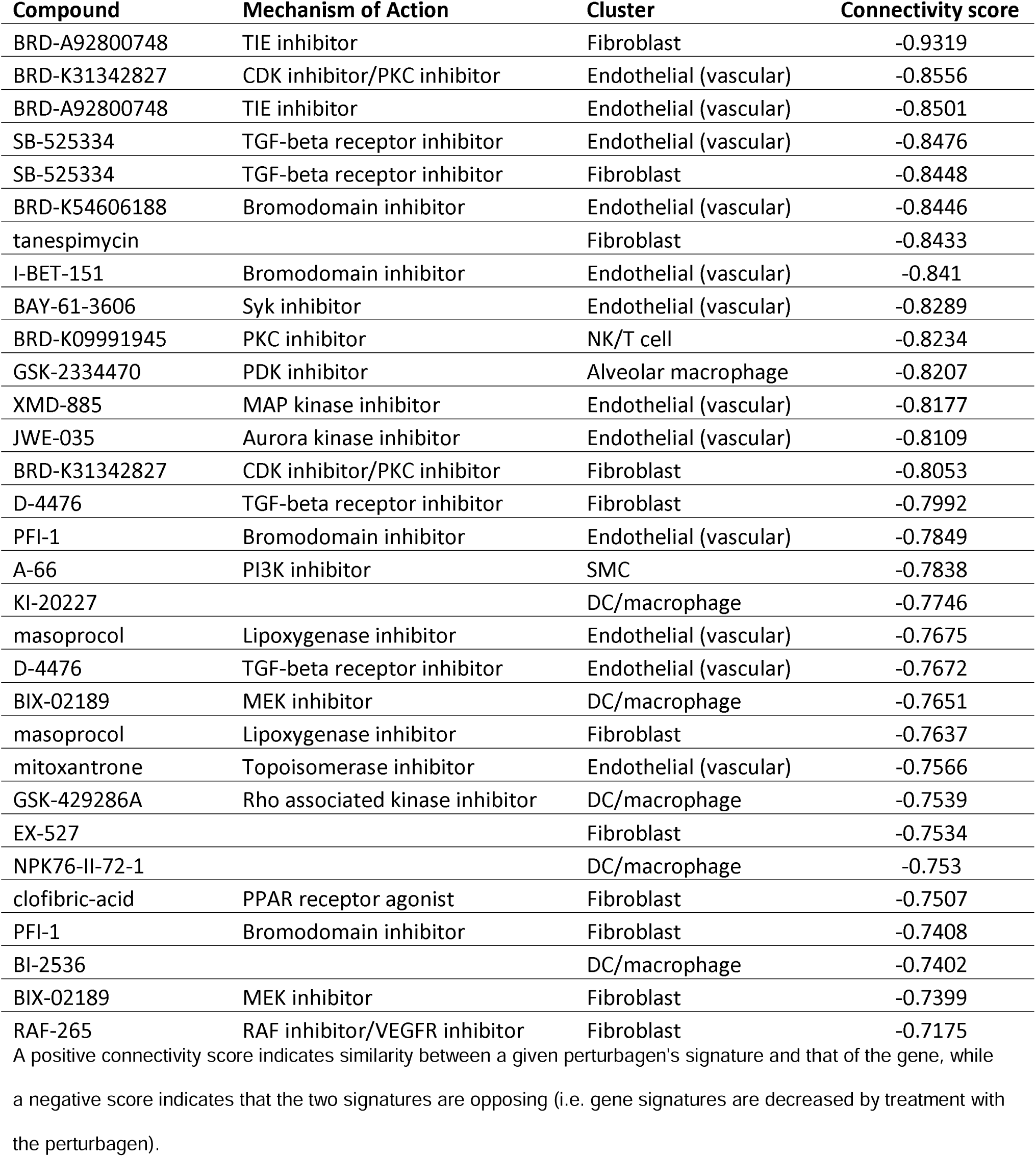
Connectivity scores and mechanisms of action for top perturbagens tested in HUVECs against SSc-PAH cell expression signatures.

Figure 6B demonstrates average connectivity scores in SSc-PAH and IPAH for phosphodiesterase inhibitors and prostanoids, drugs in common clinical use. Scores for these compounds do not meaningfully differ between the two PAH subgroups. By contrast, as shown in Figure 6C, average scores for TGF-beta receptor inhibitors, bromodomain inhibitors, TIE inhibitors, and GSK/lipoxygenase inhibitors were significantly more negative for SSc-PAH compared to IPAH (Kolmogorov-Smirnov *p-*value <0.01 for each comparison).

## Discussion

This report represents the first use of single-cell RNA sequencing of human SSc-PAH lung tissue to elucidate novel molecular targets. Two prior human PAH scRNAseq studies have examined IPAH lung tissue and isolated PAH pulmonary arteries.^45,46^ One recently published study reported single-cell gene expression patterns in SSc-PH lungs, however these specimens were taken from patients with PH generally, rather than PAH in particular, with pulmonary fibrosis contributing to PH in many cases.^47^ Ours is the only scRNAseq study to directly compare SSc-PAH versus IPAH molecular differences in the lung. Use of scRNAseq, in contrast to bulk sequencing, allows us to examine cellular diversity and disaggregate the biological processes and cell-cell signaling patterns present in these two clinically distinct PAH subtypes. We leverage this granular molecular data to produce a ranked list of compounds with potential to reverse PAH pathobiology, with a focus on SSc-PAH. Our list includes compounds that have already demonstrated efficacy in experimental PH and early-phase clinical trials and could be prioritized for further clinical investigation.

Our results show that SSc-PAH lungs differ from IPAH lungs in several respects. Fibroblasts and SMCs demonstrate a greater magnitude of differential gene expression in SSc-PAH compared to IPAH. SSc-PAH lungs exhibit an expanded fibroblast population, with a higher number of differentially expressed genes compared to IPAH fibroblasts. Similarly, there are more DEGs in SSc-PAH SMCs. In SSc-PAH versus IPAH GSEA, enrichment scores for EMT, apoptosis, myogenesis, and hypoxia are significantly higher for SSc-PAH across diverse cell types. Finally, it appears that intercellular signals target vascular endothelial cells to a greater extent in SSc-PAH compared to IPAH, particularly from ligands originating from macrophages, SMCs, and epithelial cell types.

A striking finding of our study is the extent of differential gene expression present in various epithelial cell types. Epithelial cell abnormalities were more pronounced in SSc-PAH, but differential gene expression was apparent in both SSc-PAH and IPAH epithelial cells, including in AT1 and AT2 cells and airway epithelial cells. Both IPAH and SSc-PAH AT2 cells demonstrated differential expression of genes involved in phospholipid and surfactant production. The effects of pulmonary hypertension on alveolar epithelial function are largely undefined in human PAH. However, our observations suggest altered surfactant biology and some degree of epithelial lung injury might be present.

Ciliated epithelial cells in both PAH subtypes showed upregulated *MUC5B* and *MUC5AC* relative to donor lungs, with upregulation somewhat more pronounced in SSc-PAH. Upregulated *MUC5B* is associated with increased risk of idiopathic pulmonary fibrosis and impaired mucociliary clearance in asthma,^48,49^ however *MUC5B* upregulation has not been previously reported in PAH. Other airway cell types also demonstrated differential gene expression relative to donor cells. In ligand-receptor interaction analysis, AT1 cells and basal epithelial cells played prominent roles in cell-cell signaling, with robust signaling from AT1 cells to other cell types, including vascular endothelial cells, evident in SSc-PAH in particular. The possibility of epithelial-endothelial cross-talk in PAH has been supported by a few prior studies.^50,51^ Based on these data, alveolar epithelial dysfunction and airway abnormalities might contribute, at least in part, to the pulmonary function abnormalities and diminished exercise capacity commonly observed in clinical PAH. These abnormalities could suggest novel non-vascular targets for therapeutic intervention.

Our results yield a list of drug candidates for repositioning that can be prioritized for rigorous laboratory and clinical investigation. BRD-A92800748, the top hit for SSc-PAH, is a Tie2 kinase inhibitor. Tie1 and Tie2 are angiopoietin (Ang) receptors, and the Ang/Tie pathway is dysregulated in PAH, though its exact role is controversial.^52^ Tie2 expression is increased at the mRNA and protein level in PAH lung tissue and potentiates SMC hyperplasia.^53^ However, in different experiments, both Ang-Tie2 potentiation and Ang-Tie2 inhibition have been found to improve experimental PH,^52,54^ which likely reflects the complexity of a signaling system exerting varying effects on angiogenesis and vascular integrity and function. Notably, Tie2 is an important regulator of peripheral vasculopathy in SSc.^55^

Other top compounds on our list have strong preclinical and/or clinical evidence in support of efficacy in PAH. Several GSK-3 inhibitors strongly opposed our SSc-PAH expression signatures. GSK-3 contributes to PAH pathobiology via myofibroblast activation and beta-catenin signaling, and inhibiting this pathway can prevent PH and RV remodeling in animal models.^56–58^ Bromodomain inhibitors, which our data suggest could be more efficacious in SSc-PAH, inhibit BET proteins, epigenetic regulators of transcriptional programs that regulate cell proliferation and survival. Bromodomain inhibitors reverse endothelial and SMC abnormalities *in vitro* and pulmonary hypertension *in vivo* across multiple rodent models of PH.^59^ A small single-arm pilot study of the bromodomain inhibitor apabetalone demonstrated reductions in PVR and improvement in cardiac function in seven patients,^60^ and a phase II clinical trial is planned (NCT04915300). Our data suggest TGF-beta receptor inhibitors, which include the drug sotatercept, could also be more efficacious in SSc-PAH.

There are important limitations to this study. Unlike prior scRNAseq studies of fresh lung tissue, to directly compare equal numbers of IPAH and SSc-PAH specimens, this study utilized banked tissues. Consequently, our cell count and sequencing depth are somewhat lower than in prior studies. Nevertheless, the sequencing depth was sufficient to yield feature counts typical of 10X Genomics platforms, and RNA was of sufficiently quality for downstream analyses. We were well-powered to detect expression differences between major cell types, but we were not able to sub-cluster cell types or detect rare cell populations. This limitation reflects a tradeoff whereby banking tissues makes possible accumulation of specific phenotypes of interest (e.g., SSc-PAH), but can lead to reduced data resolution. Our samples were from end-stage PAH patients. Some gene expression patterns may therefore reflect compensatory processes or were induced by treatment with PAH-specific therapies. However, this is the profile of patients most in need of additional, novel therapies. Finally, the assumption that mRNA levels correlate with protein expression is not always valid. However, at present, scRNAseq is a far more mature technology than single-cell proteomic profiling.

In conclusion, here we demonstrate an array of targetable molecular differences in human SSc-PAH versus IPAH lung tissues. Some of these SSc-specific molecular aberrations might underlie observed disparities in SSc-PAH epidemiology. Future studies focused on novel therapeutic targets with the potential to improve outcomes in SSc-PAH are warranted.

## Acknowledgements

C.E.S. had full access to all the data in the study and takes responsibility for the integrity of the data and accuracy of the data analysis. T.T. contributed substantially to the study design, data analysis and interpretation, and the writing of the manuscript. J.C.C., T.G., M.P., C.C., S.E.N, S.L.K, D.T.R., M.P.C. M.E., T.M.K., S.C.M., R.L.D., P.M.H., L.A.S., K.S., and M.A.A. contributed substantially to interpretation of the data for work and revision of the manuscript. Tissue samples were provided by the Pulmonary Hypertension Breakthrough Initiative (PHBI).

