## Supplemental Figures for "Single-cell transcriptomics reveal diverging pathobiology and opportunities for precision targeting in scleroderma-associated versus idiopathic pulmonary arterial hypertension"

### Supplemental Methods

*Tissue handling and processing*

We identified twelve banked lung specimens from the Pulmonary Hypertension Breakthrough Initiative (PHBI) tissue bank for sequencing: four SSc-PAH lung specimens, four IPAH lung specimens, and four failed donor specimens judged suitable to serve as controls. Classification of PAH subtype was adjudicated by expert agreement. PHBI identifiers and clinical variables associated with the samples selected are listed in Table 1. Twenty-five uM tissue sections were cut from banked tissue blocks. Peripheral lung samples were prioritized to capture distal pulmonary vessels. Tissue sections were rehydrated and transferred to a 1.5ml tube. Tissues were de-paraffinized using a series of sequential xylene, ethanol, and PBS washes. Pestle dissociation was performed using a dissociation enzyme mix prepared per manufacturer's instructions. Following dissociation, tissues were centrifugated and tissue pellet was resuspended, after which cell concentration was determined. Single cells were isolated following manufacturer's protocols designed to maximize RNA integrity and minimize degradation.

*Probe Hybridization*

Single-cell RNA sequencing (scRNAseq) was performed using the 10X Genomics Chromium Flex platform, which is a droplet-based technology optimized for formalin-fixed paraffin-embedded (FFPE) tissues. Whole transcriptome probe pairs were added to the fixed sample. Together, probe pairs hybridize to their complementary target RNA in an overnight incubation.

*GEM Generation and Barcoding*

Single cells were encapsulated into droplets in the presence of gel beads in-emulsion (GEMs) carrying barcoded oligonucleotides. Samples containing different Probe Barcodes were pooled together and the unbound probes were washed off. After pooling and washing the samples, GEMs were generated by combining barcoded Gel Beads, a Master Mix containing pooled cells, and Partitioning Oil onto Chromium Next GEM Chip Q. A pool of GEM Barcodes was sampled separately to index the contents of each partition. Within these droplets, reverse transcription occurs, converting RNA into cDNA while simultaneously attaching a unique barcode and UMI (Unique Molecular Identifier) to each cDNA molecule. Following GEM generation, the gel bead was dissolved releasing barcoded primers, while any co-partitioned cell was lysed. Following barcoding, the GEMs are broken, and ligated products are pre-amplified, then cleaned.

*Library construction and sequencing*

The cDNA then underwent library construction steps, including fragmentation, end-repair, A-tailing, adapter ligation, and PCR amplification. Library quality and quantity were assessed using the Agilent Bioanalyzer system and Qubit fluorometer, respectively, before proceeding to sequencing. Sequencing was carried out on an Illumina NovaSeq platform. The sequencing data were processed and demultiplexed using 10X Genomics Cell Ranger software, which aligned reads to the reference genome.

*Statistical analysis*

Number of transcripts per cell and percent mitochondrial genes per sample were assessed for quality control. RNA barcodes were filtered on total UMI count (> 500 UMIs), feature count (> 250 features), and percentage of mitochondrial genes (< 5%). Following quality control procedures, including batch correction performed with Harmony,[1] data normalization and principal components analysis was performed using Seurat v4.3.0.[2] Leiden clustering was performed on the dimensionally-reduced data. Clusters were identified using marker genes via the FindMarkers function in Seurat and differential expression comparing each cluster to all other cells in the dataset. Significance of differential expression was tested using the Mann-Whitney U test. Clusters were manually annotated using canonical cell type markers then compared with annotated publicly available single-cell expression data using LungMap.[3] A Benjamini-Hochberg correction was performed to account for multiple testing. Genes were considered significantly differentially expressed if the adjusted p-value was <0.05.

Gene set enrichment analysis (GSEA) was performed using Hallmark gene sets from the Molecular Signatures Database (MSigDB).[4] The GSEA algorithm fgsea v1.24.0 evaluates the distribution of predefined gene sets within the ranked list of all genes from differential expression analysis, thereby identifying gene sets that are overrepresented at the top or bottom of this list.

To infer differential transcription factor (TF) activity across disease subtypes, we utilized UCell, an analytical tool developed for quantifying the activity of transcription factors and gene sets in single cells.[5] The UCell algorithm scores each cell based on the expression of a pre-defined set of target genes associated with each TF, using a rank-based metric that accounts for both the presence and relative expression level of these target genes. For this analysis, target gene sets for each TF were curated from MSigDB. After scoring individual cells, we performed differential expression analysis on these scores, stratified by cell cluster.

CellPhoneDB was used to identify pairs of potential receptors and ligands to infer intercellular signaling among SSc-PAH and IPAH cells.[6] NicheNet's protein-protein interaction network was used to identify potential receptors upstream of transcription factors in known protein-protein signaling networks.[7] Scores for possible signaling pathways between clusters of cells were calculated as the multiple of the z-scored ligand expression, receptor expression, and transcription factor activity scores generated via UCell.[8] The sum of inter-cluster signaling scores was then taken as the sum of the highest scores, limited to one score per transcription factor. ComplexHeatmap v2.14.0 and circlize v0.4.15 were used for visualizations.[9, 10]

We leveraged the Connectivity Map (CMap) platform[11, 12] to perform pharmacotranscriptomic screening using cluster-specific DEGs from SSc-PAH vs. controls and IPAH vs. controls as input signatures. CMap utilizes a pattern-matching algorithm to compare input gene signatures against its database of reference expression profiles generated by treating various cell lines with thousands of perturbagens, including small-molecule compounds. For each input signature, CMap generates connectivity scores for each screened compound in the database, which represents the degree of similarity or opposition between treatment-induced gene expression changes and a specific input signature. A negative connectivity score indicates the potential for a given perturbagen to reverse the inputted gene expression pattern.

**Supplemental Figure Legends**

**Supplemental Figure 1a and 1b: Sample Quality Control** Distributions of features detected (genes per cell on the y-axis) by UMI count (on the x-axis) in scRNA-seq data from lung tissue samples. Each point represents an individual cell, with colors indicating percentage of mitochondrial DNA.

**Supplemental Figure 2: Batch Correction** UMAP plot of scRNA-seq data before (top) and after (bottom) batch correction using Harmony. Cells are colored by sample source and grouped by cell type cluster (after correction), demonstrating the effectiveness of batch correction in integrating data across different samples.

**Supplemental Figure 3a and 3b: Cell Type Proportions by Condition** Bar plot showing the proportion of each identified cell type across different conditions (IPAH, SSc-PAH, and donors). Colors represent different conditions. P-values are shown above the brackets comparing across different conditions.

**Supplemental Figure 4a-11b: Differential Gene Expression by Cell Type** Volcano plots of differentially expressed genes between SSc-PAH and donor samples, IPAH and donor samples, and SSc-PAH versus IPAH are shown for each cell type with genes demonstrating highly significant differential expression. Significant genes (adjusted p-value < 1E-4) are highlighted, showing both upregulated (to the right of the vertical bar) and downregulated (to the left of the vertical bar) genes. For each volcano plot, the x-axis is the log scaled fold-change between the two groups of cells being compared. The y-axis is the negative log adjusted p-value. The horizontal dashed line represents an adjusted negative log p-value of 0.05. Beneath the volcano plots appear running enrichment plots, from gene set enrichment analysis (GSEA), for the gene sets with the normalized enrichment scores of greatest magnitude. For a given gene set, differentially expressed genes are ordered along the x axis by significance, with the most upregulated on the left and the most downregulated on the right; each tick mark represents a gene in the set. If the gene set is highly upregulated, tick marks aggregate to the left side of the plot, and visa-versa for downregulated gene sets. The blue line shows the running enrichment score, a score indicating whether the gene set is showing up more or less often than would be expected due to chance alone at the beginning or the end of the complete set of genes. The peak of the blue line (either positive or negative) represents the enrichment score. Each grey line represents one random permutation of the data, of which there are 100 plotted.

**Supplemental Figure 13a and 13b: Common Differential Expression in SSc-PAH and IPAH** Scatter plots showing the average log2 fold change (log2FC) of differentially expressed genes (DEGs) in SSc-PAH compared to IPAH across cell clusters. Each point represents a gene, with its position on the plot indicating the average log2FC in SSc-PAH (y-axis) versus IPAH (x-axis). The blue dashed line represents the line of equality (y = x), where genes would fall if they had equal expression changes in both conditions. Points above the line are upregulated in SSc-PAH relative to IPAH, while points below the line are downregulated in SSc-PAH relative to IPAH.

**Supplemental Figure 14: Topologic Overview of Transcription Factor Activity Differences by UCell** Jitter plots showing the significance of differences in transcription factor (TF) activity across various cell clusters between IPAH and control samples (top) and SSc-PAH and control samples (bottom). Each point represents a transcription factor, with its position on the plot indicating the −log10(p-value) of the difference in activity. The y-axis represents the significance of the differential TF activity, while the x-axis lists the different cell clusters. Points colored differently indicate statistically significant differences in TF activity between the compared conditions, with higher values on the y-axis representing greater significance. Gray dots represent TFs without statistically significant differences in activity between conditions.

**Supplemental Figure 15: AT1 Signaling to Vascular Endothelial Cells** At top, heatmaps show quantitative scores for transcription factor (TF)-ligand-receptor interactions in IPAH (left) and SSc-PAH (right) samples. The heatmaps display significant signaling scores for various interactions, with the color intensity representing the interaction strength (red indicating higher scores). Below, UMAP plots illustrate the cluster-specific expression patterns of specific molecules involved in the highlighted interactions across different cell types. The plot highlights TGFβ2_TGFβ receptor1/2, TGFβ2_TGFβ receptor1, and IL1β_IL1 receptor expression in IPAH, and VEGFA_FLT1 complex, CLEC4N_CNTNAP1, ACTH_MC1 receptor, ANXA1_FPR2 receptor, and ALB_FcRn complex in SSc-PAH.

**Supplemental Figure 16: SMC Signaling to Vascular Endothelial Cells** At top, heatmaps show quantitative scores for transcription factor (TF)-ligand-receptor interactions in IPAH (left) and SSc-PAH (right) samples. The heatmaps display significant signaling scores for various interactions, with the color intensity representing the interaction strength (red indicating higher scores). Below, UMAP plots illustrate the cluster-specific expression patterns of specific molecules involved in the highlighted interactions across different cell types. The plot highlights the PGF-FLT1 complex and TGF-beta signaling pathways

**Supplemental Figure 17: Alveolar Macrophage Signaling to Vascular Endothelial Cells** At top, heatmaps show quantitative scores for transcription factor (TF)-ligand-receptor interactions in IPAH (left) and SSc-PAH (right) samples. The heatmaps display significant signaling scores for various interactions, with the color intensity representing the interaction strength (red indicating higher scores). Below, UMAP plots illustrate the cluster-specific expression patterns of specific molecules involved in the highlighted interactions across different cell types. The plot highlights VEGFB-FLT1 and TGFB1-TGF-beta receptor interactions.

**Supplemental Figure 18** Chord diagrams showing cell-cell signaling differences highlighting VEGF, TGF-beta, and PDGF signaling between IPAH and SSc-PAH for fibroblasts, alveolar macrophages, and dendritic cells/macrophages. Each segment around the circle corresponds to a different cell cluster identified in the dataset. The length of colored segments indicates the relative abundance of each cell type. Chords connecting the segments represent ligand-receptor interactions, with the thickness of each chord proportional to the interaction strength. Colors are used to distinguish between different cell types and their interactions. Arrowheads indicate directionality from ligand to receptor.

**Supplemental Figure 19a and 19b** Violin plots showing the expression patterns for common signaling pathways implicated by the data, including of VEGFA, VEGFB, FLT1, TGFB1, PDGFB, and PDGFR. Expression level is on the y-axis, and different cell types are listed along the x-axis colored by failed donor (yellow), IPAH (blue), and SSc-PAH (green) samples.

**Supplemental Figure 20: Pharmacotranscriptomic Screening Overview** A) Heatmaps of connectivity scores for individual compounds tested against IPAH (left) and SSc-PAH (right) gene expression signatures. Individual compound names are on the y-axis at left, with the corresponding mechanism of action (if known) listed at right; cell types are on the x-axis. Scores are depicted for compounds with FDR-corrected p-value <0.05. Dot color corresponds to connectivity score, with dark purple representing -1, and light blue representing -0.7. Dot size corresponds to -log10(p-value). Similar heatmaps showing average scores when compounds with mechanisms of action in common are grouped are shown for IPAH in (B) and SSc-PAH in (C).

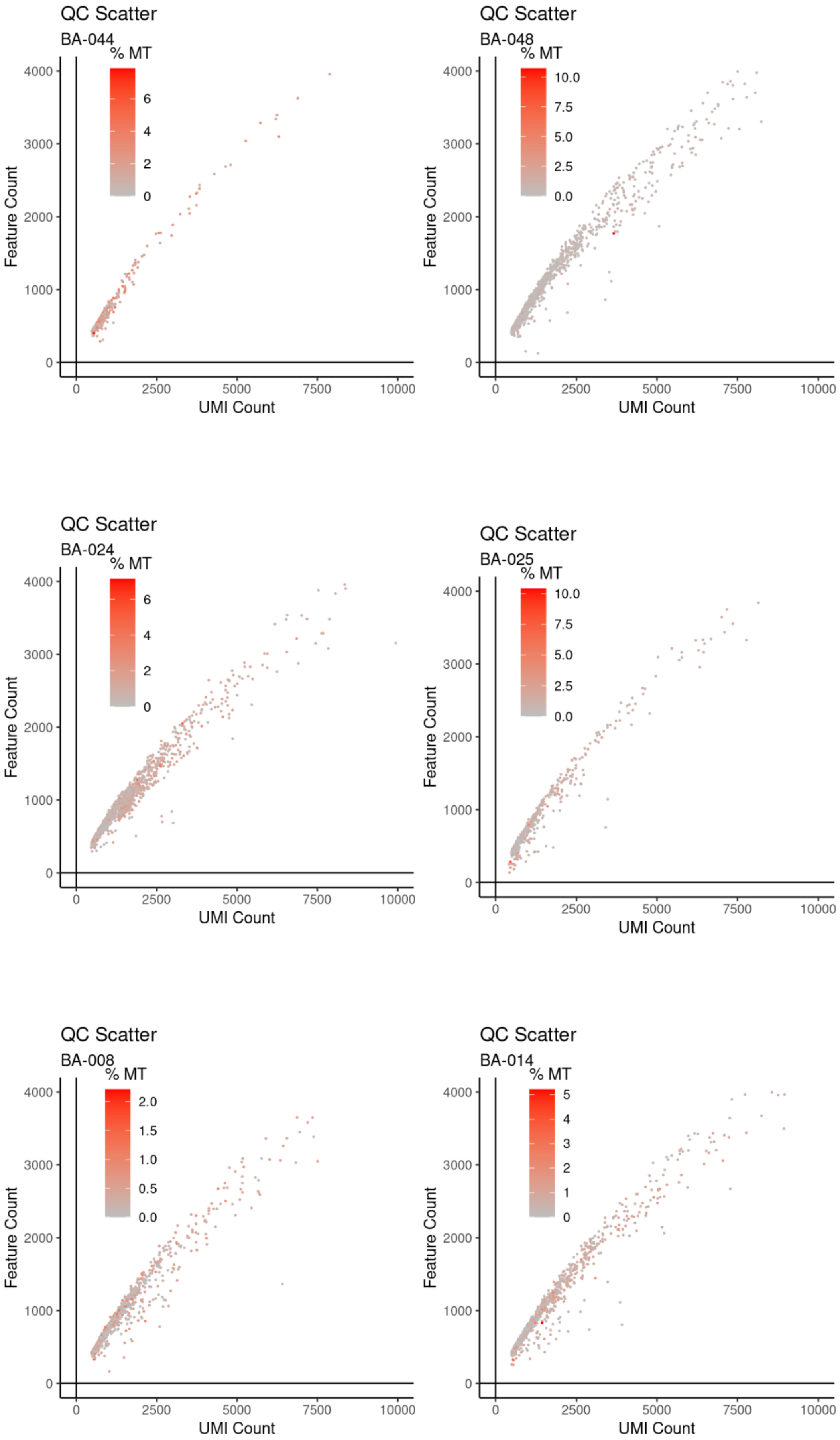


Supplemental Figure 1a

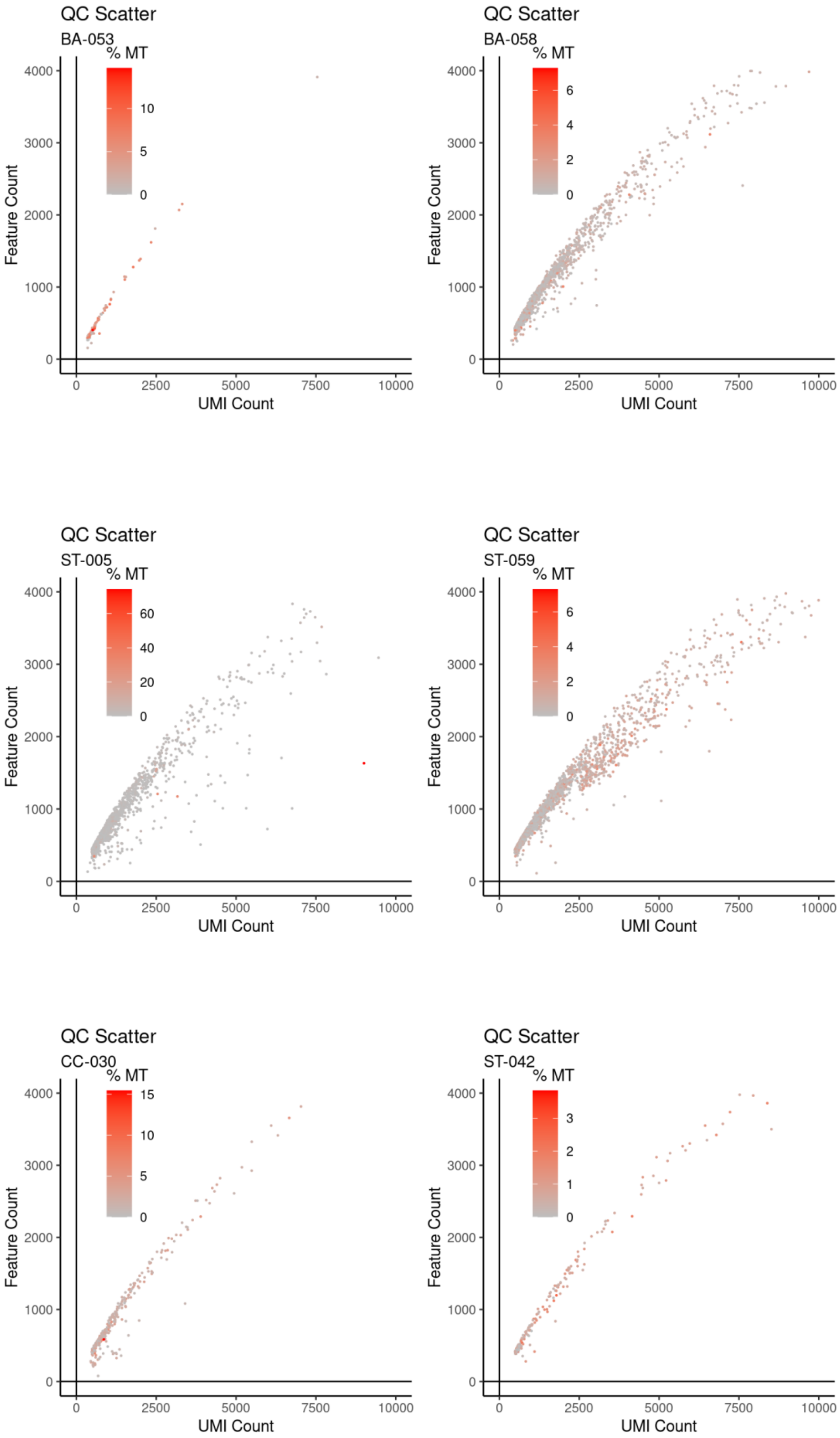


Supplemental Figure 1b

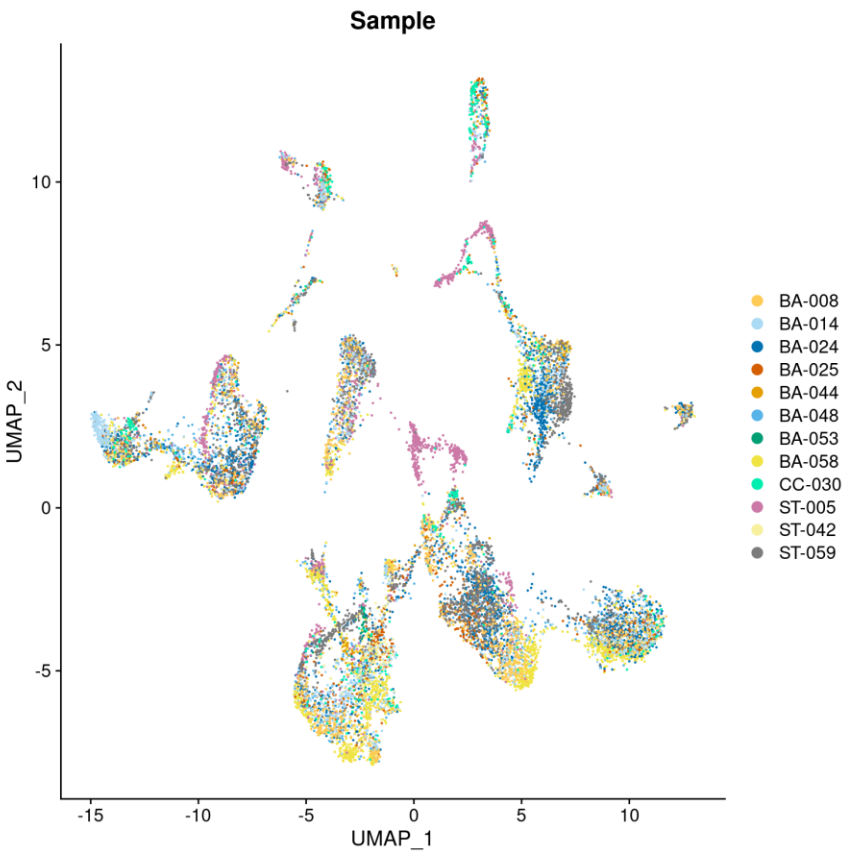


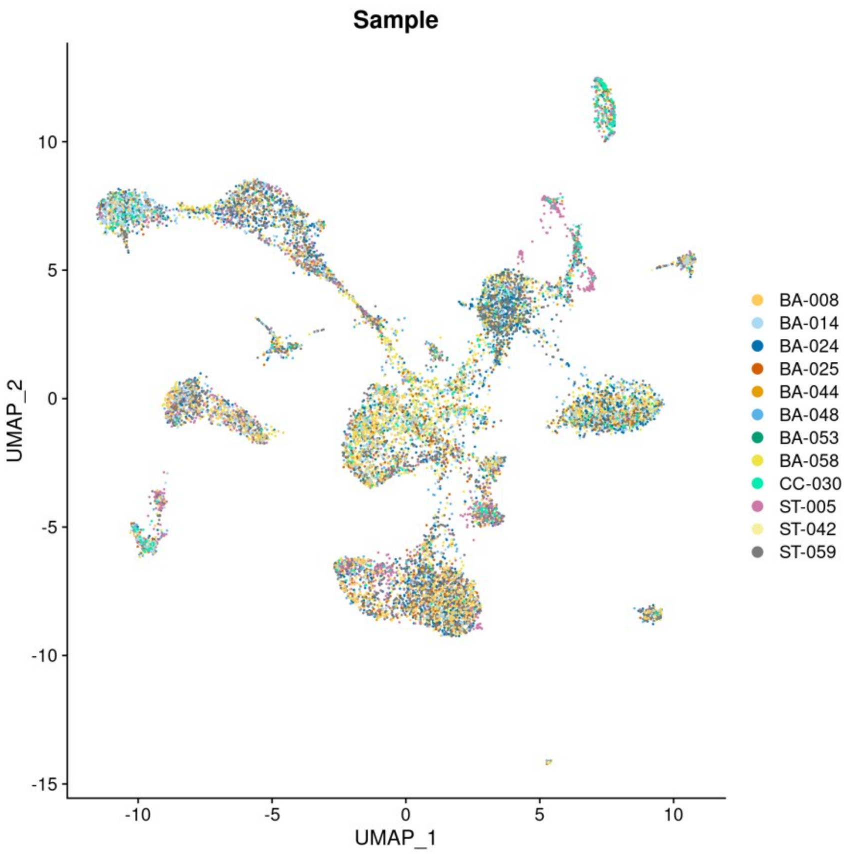


Supplemental Figure 2

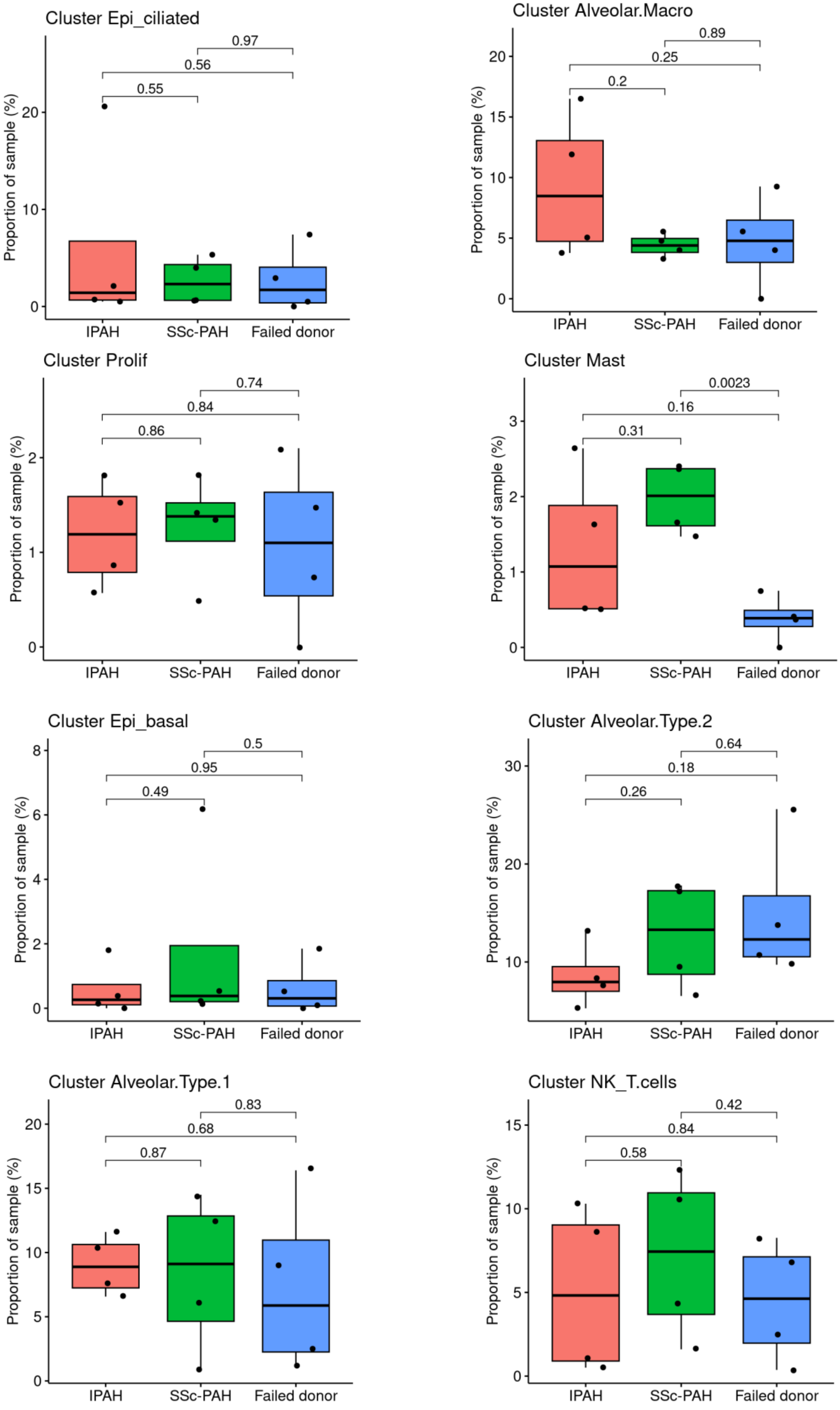


Supplemental Figure 3a

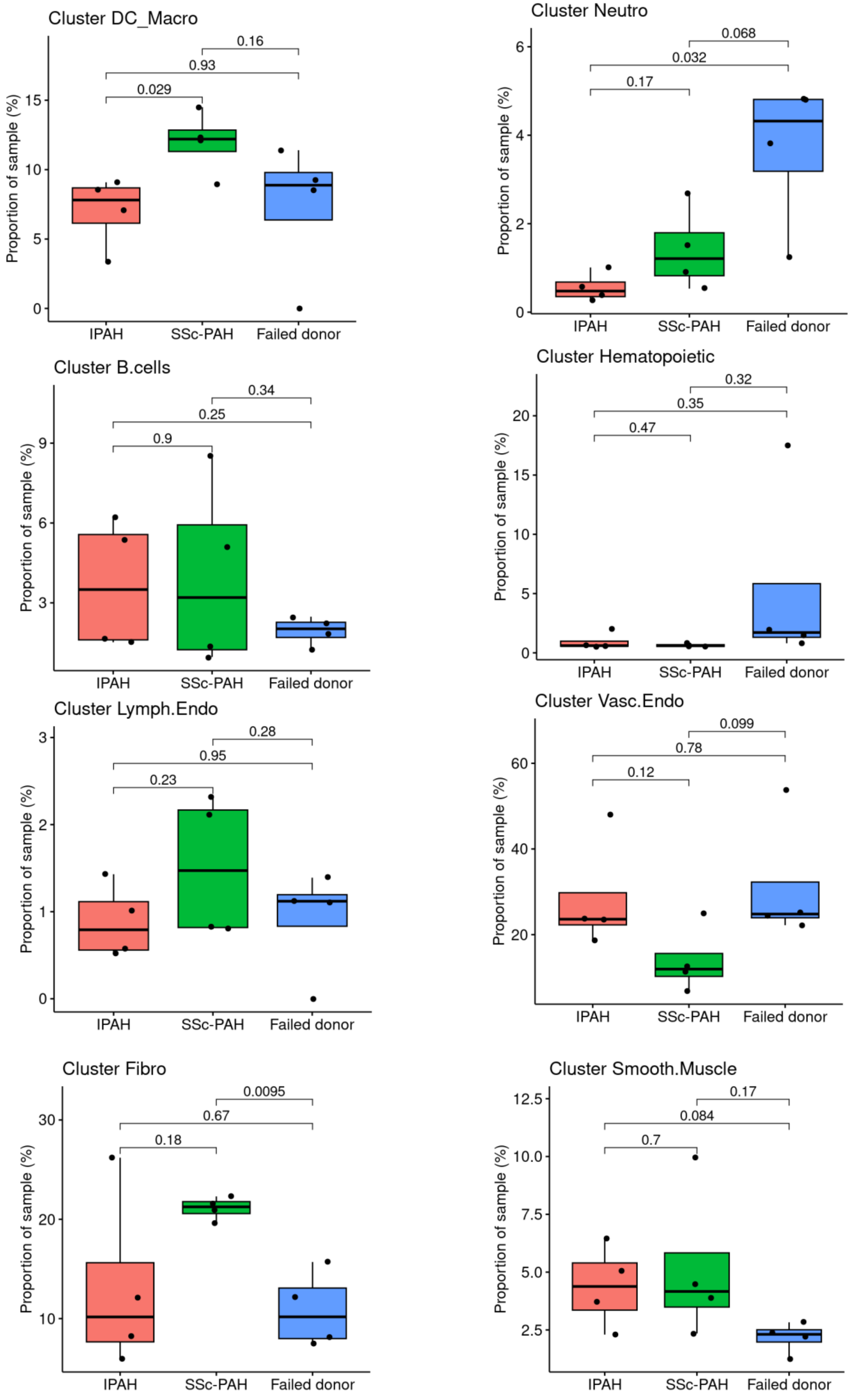


Supplemental Figure 3b

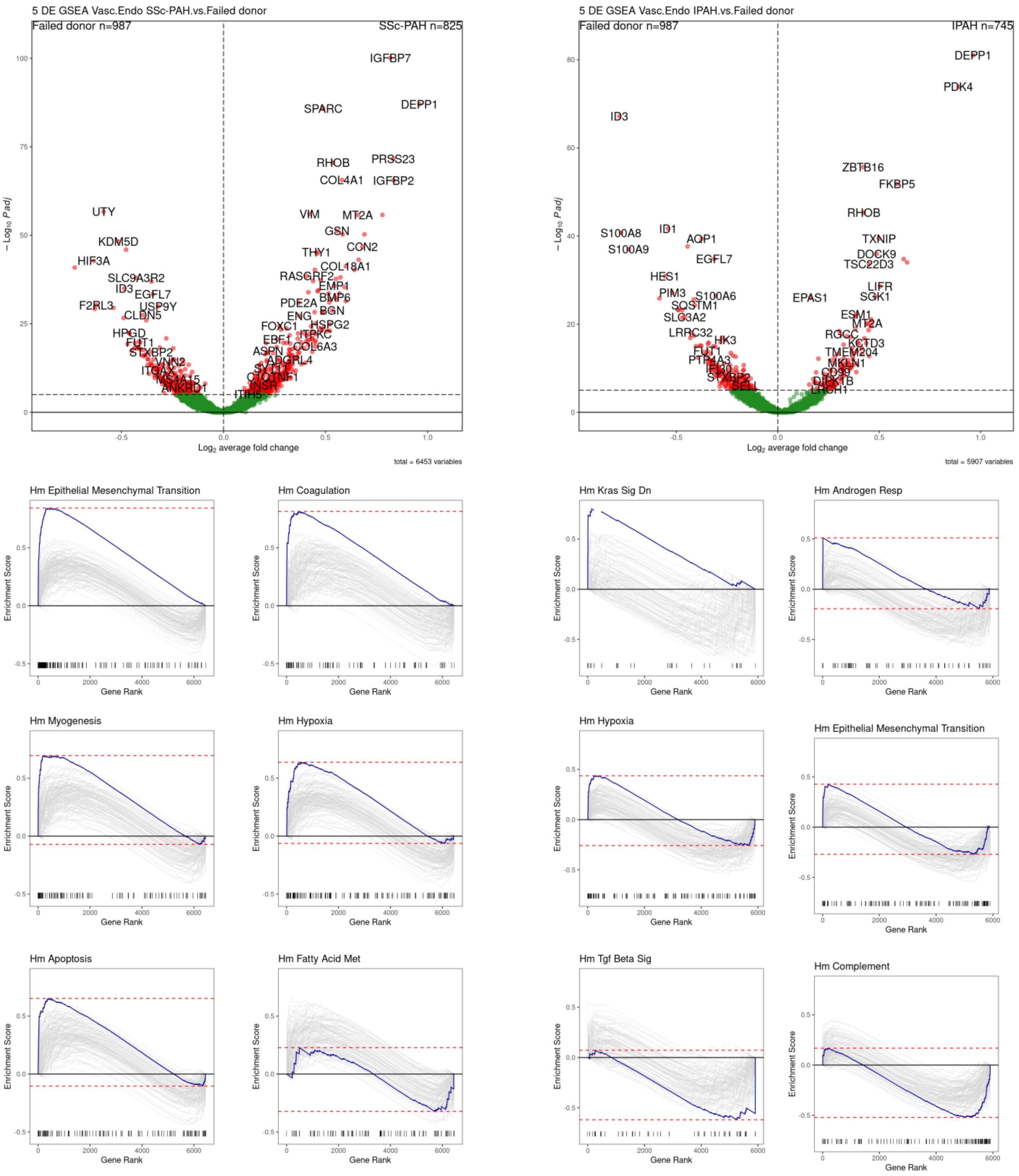


Supplemental Figure 4a

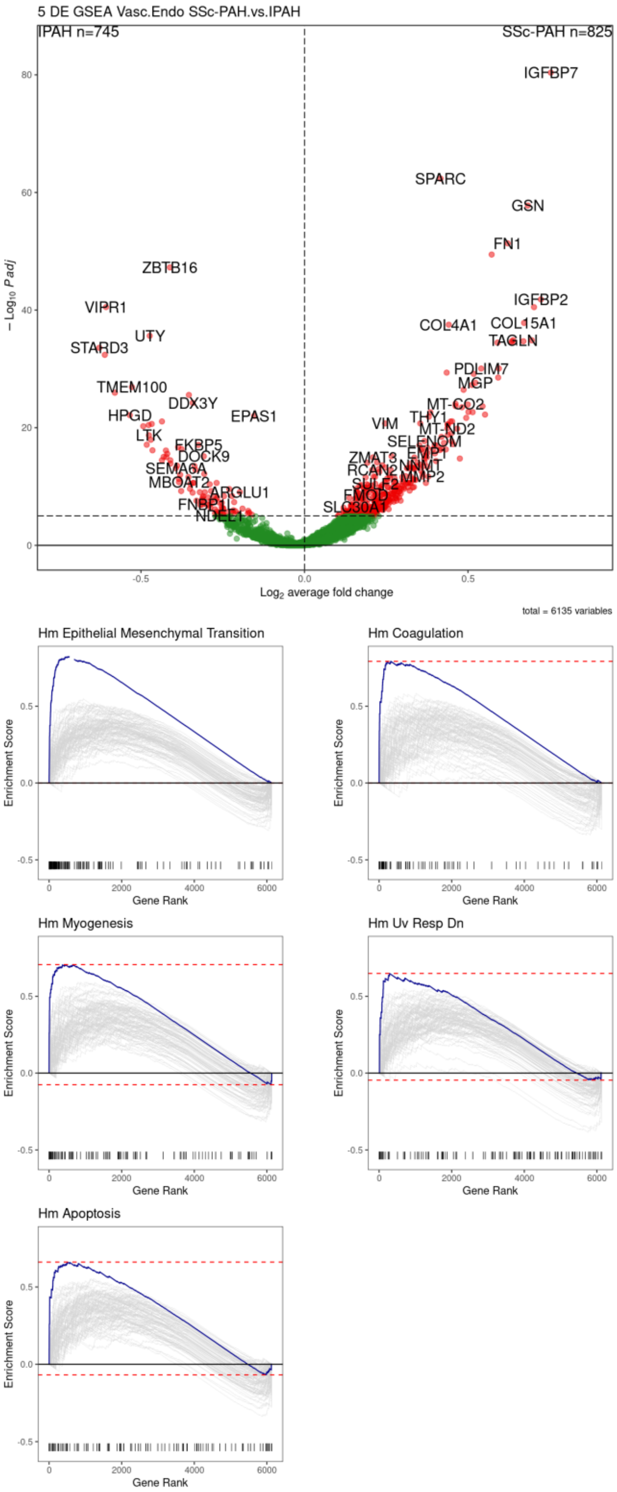


Supplemental Figure 4b

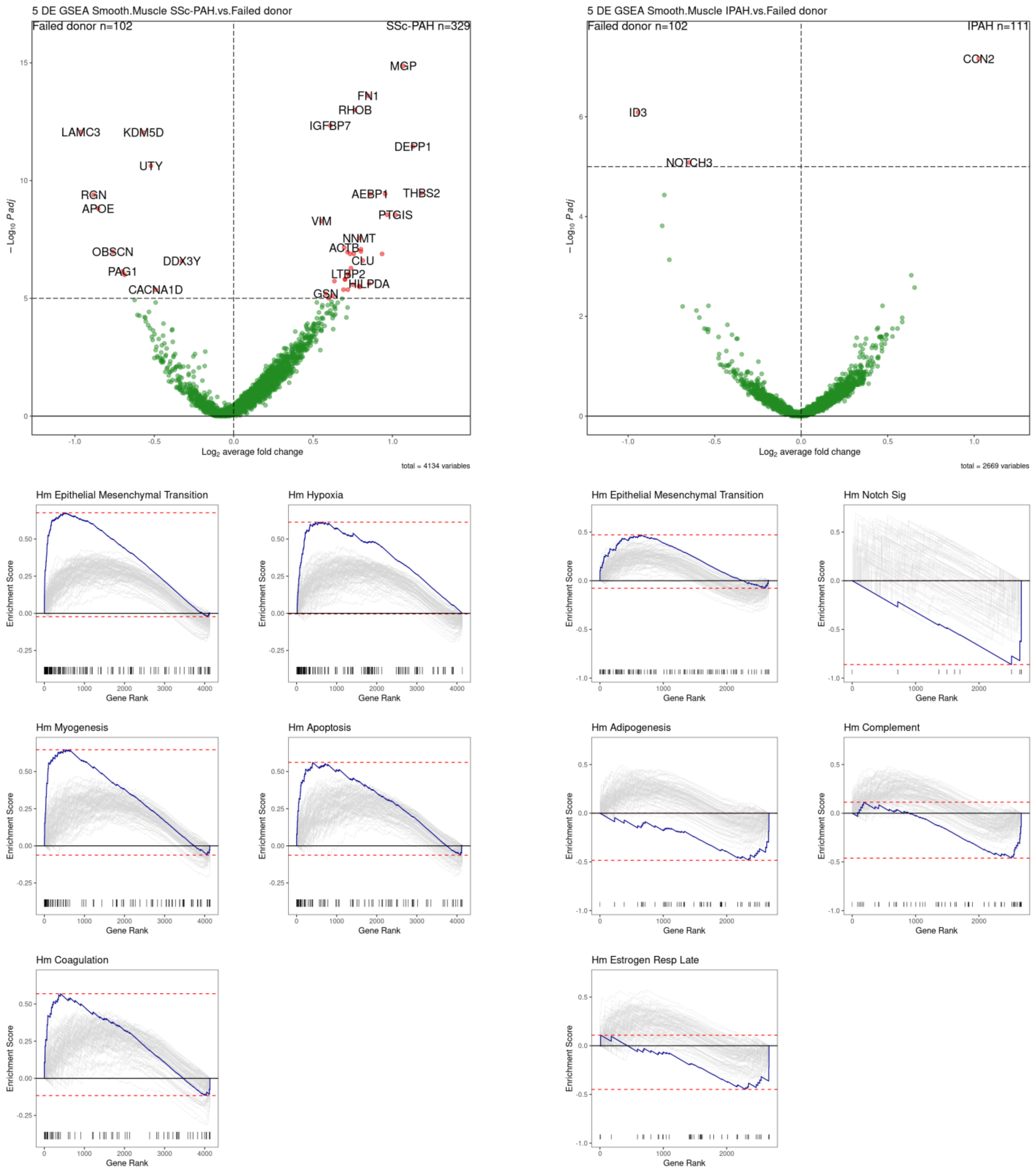


Supplemental Figure 5a

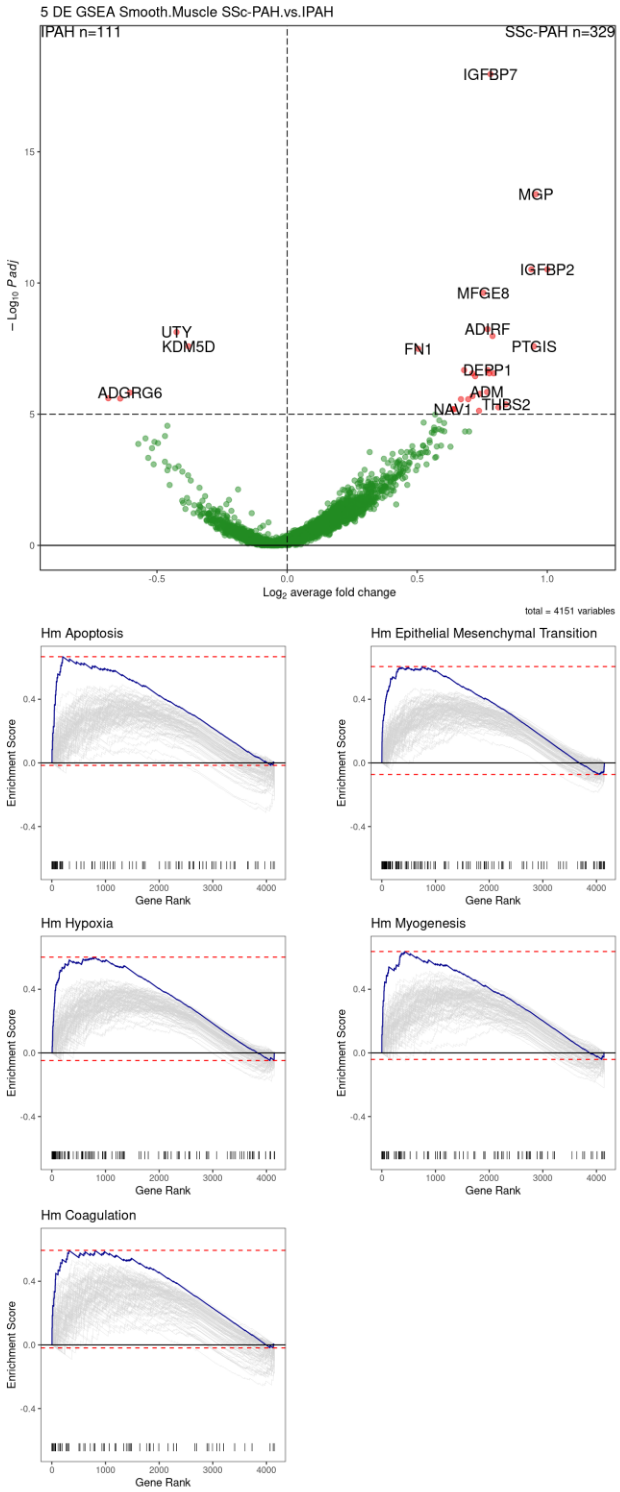


Supplemental Figure 5b

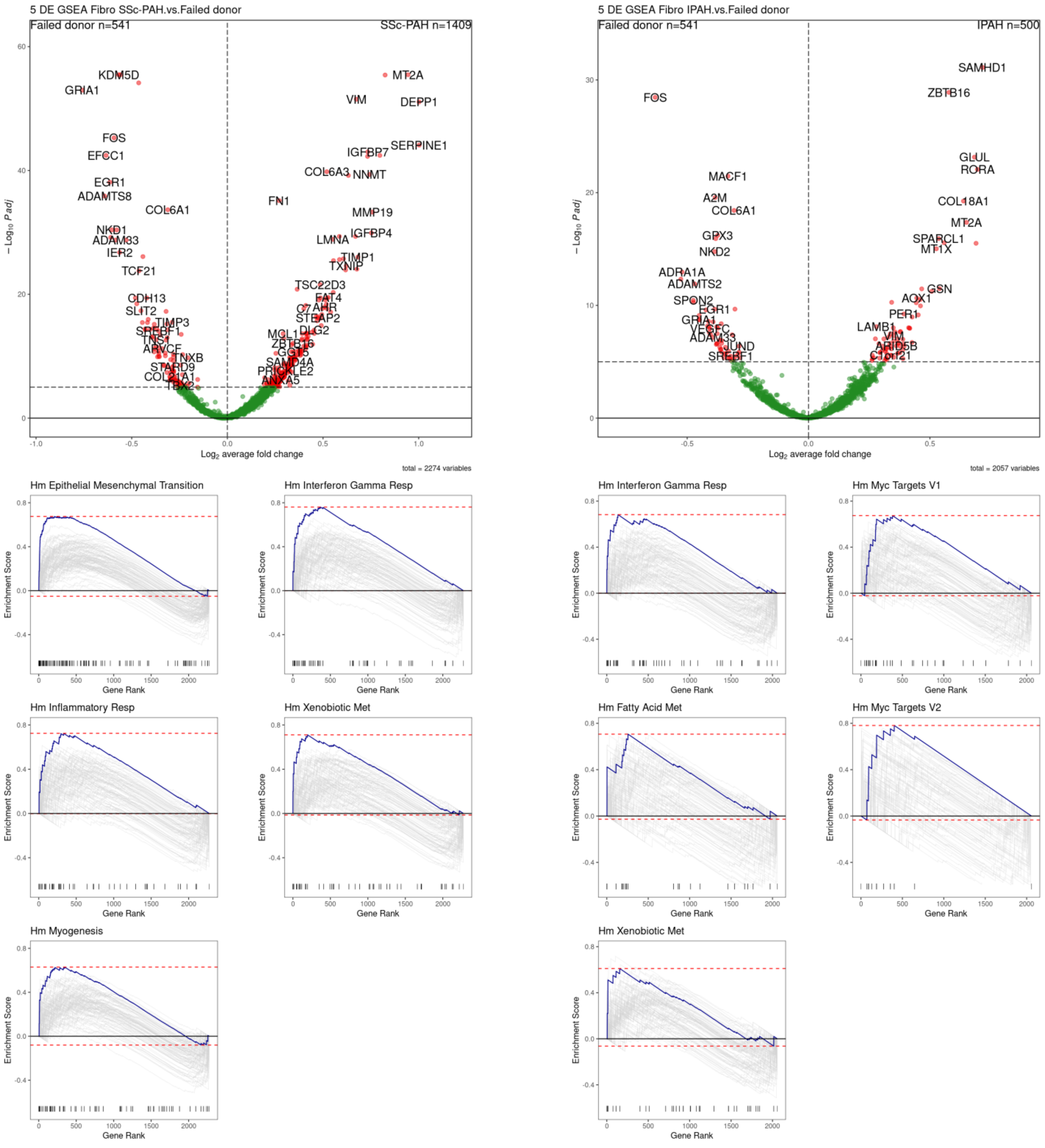


Supplemental Figure 6a

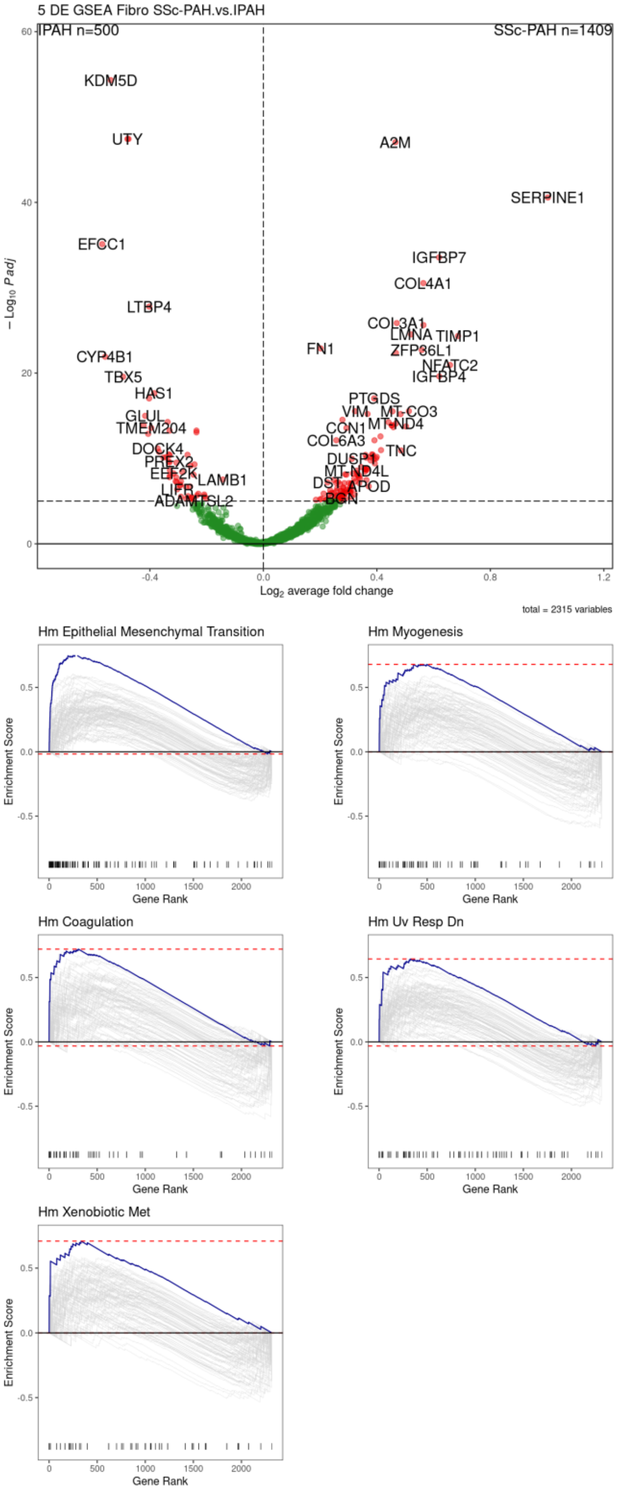


Supplemental Figure 6b

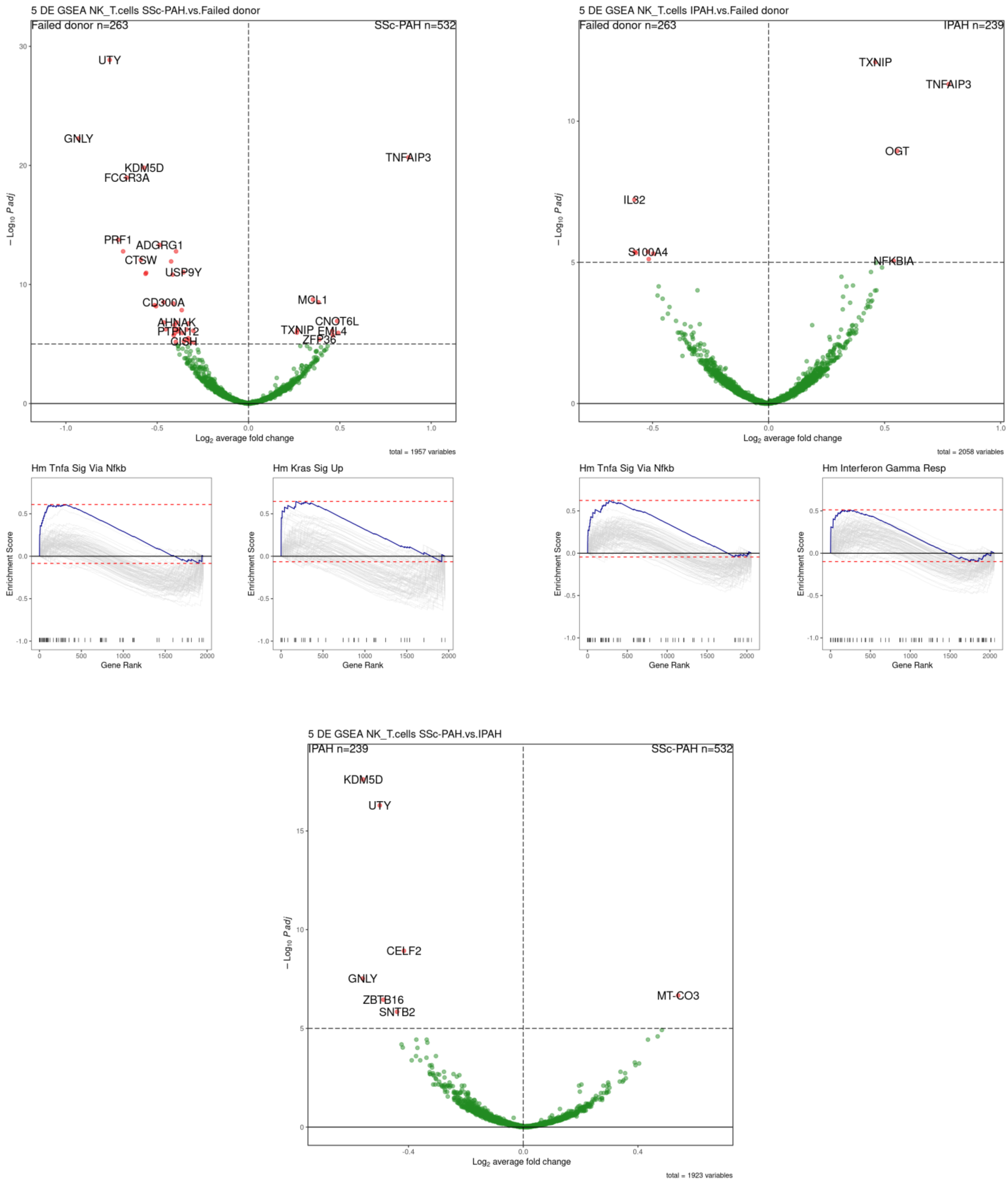


Supplemental Figure 7

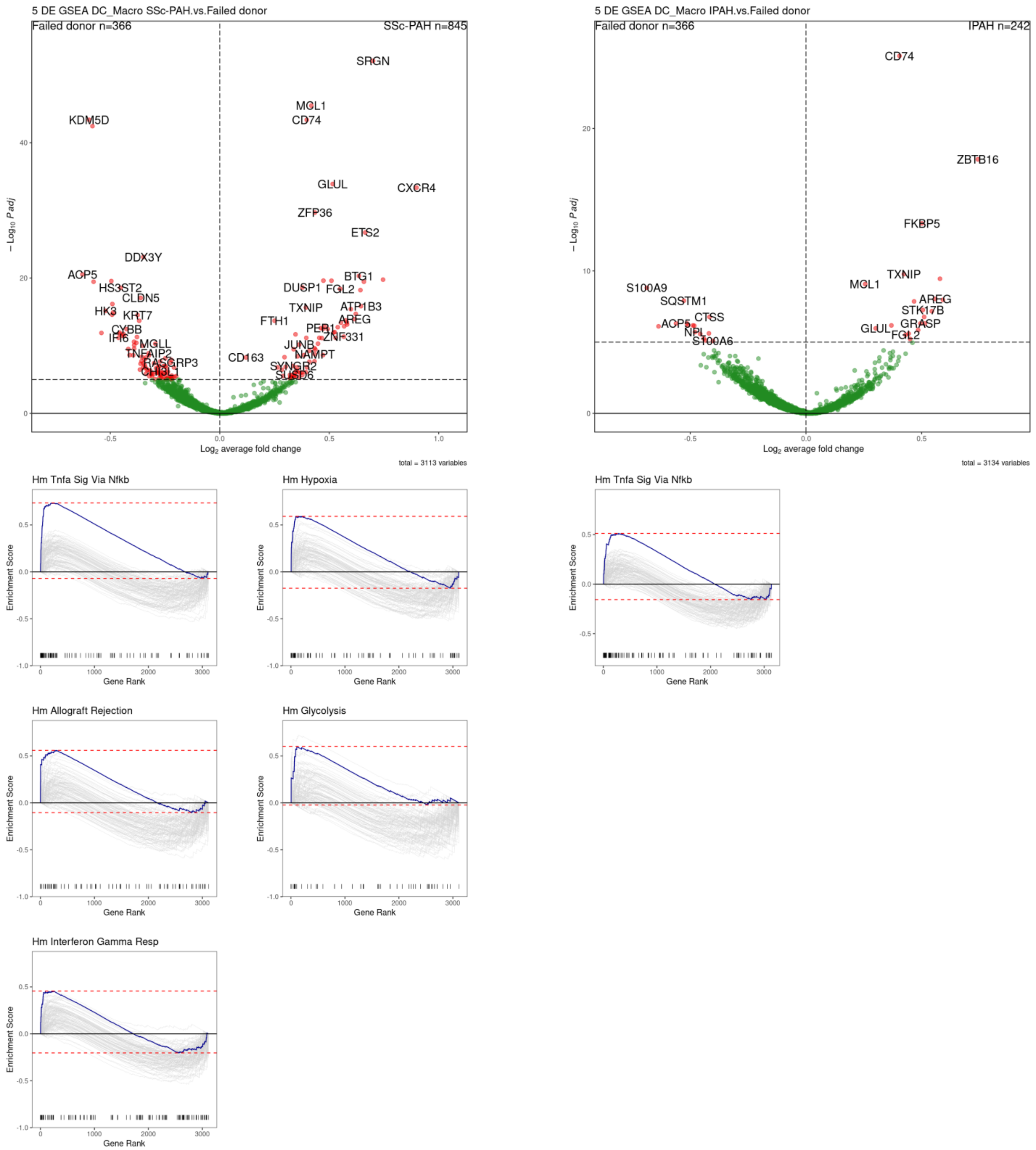


Supplemental Figure 8a

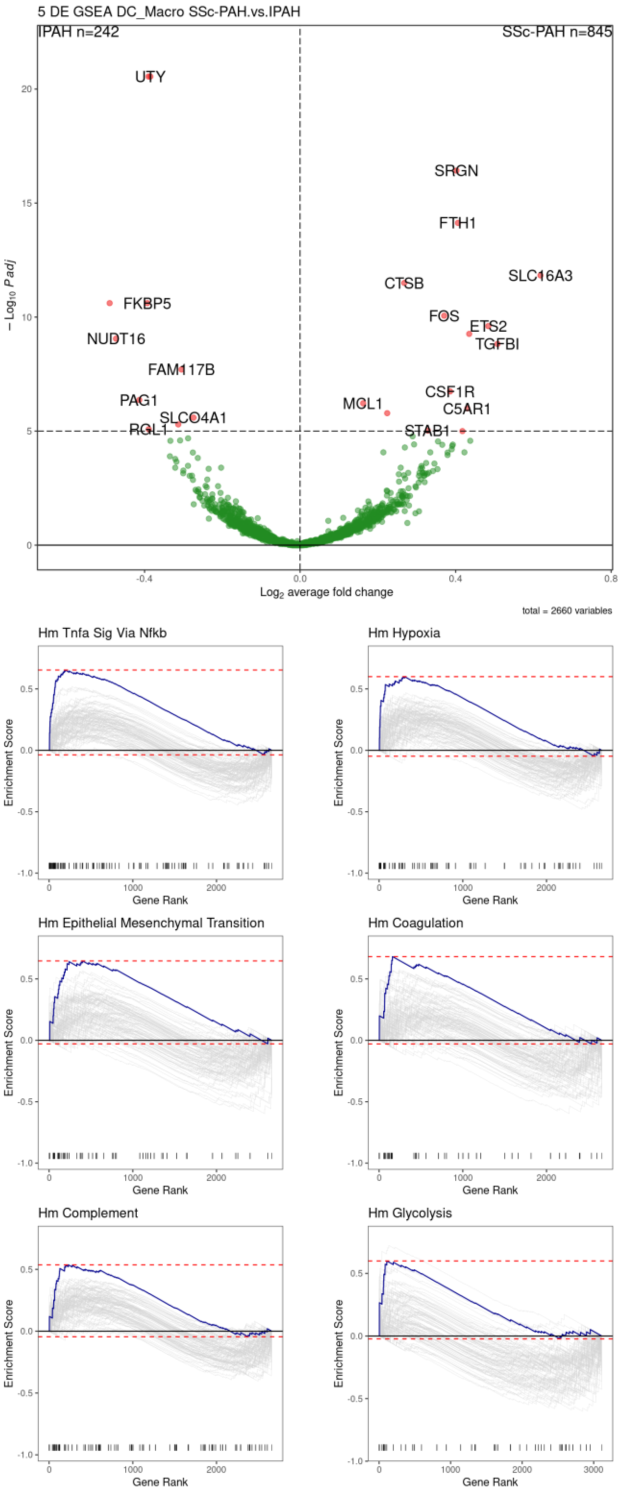


Supplemental Figure 8b

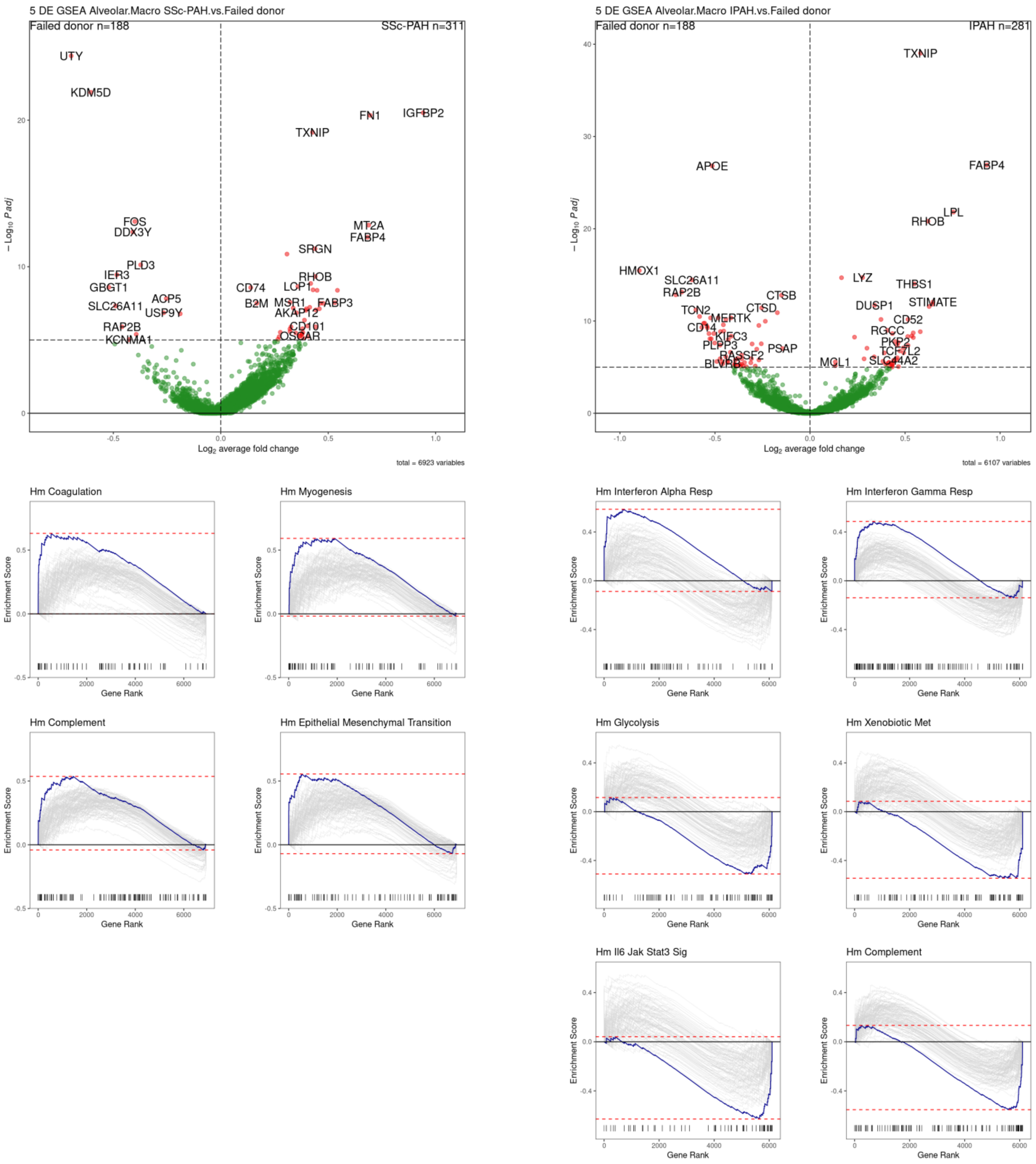


Supplemental Figure 9a

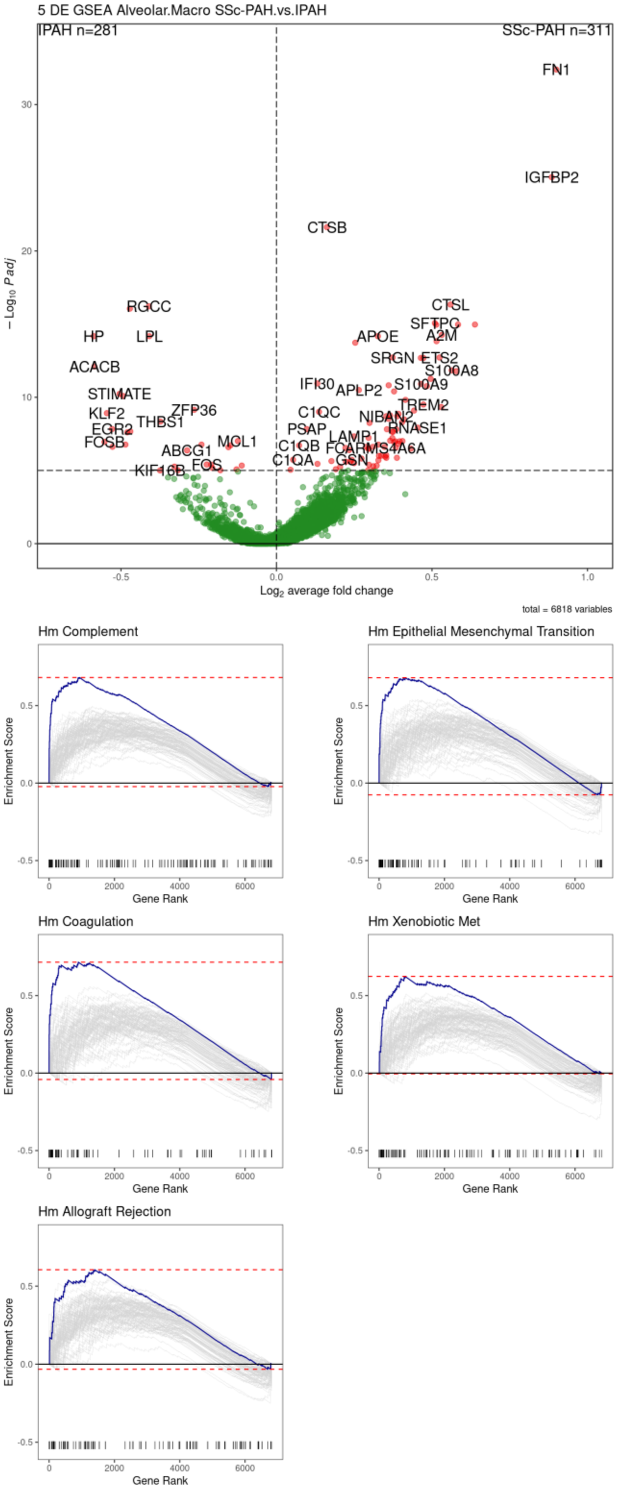


Supplemental Figure 9b

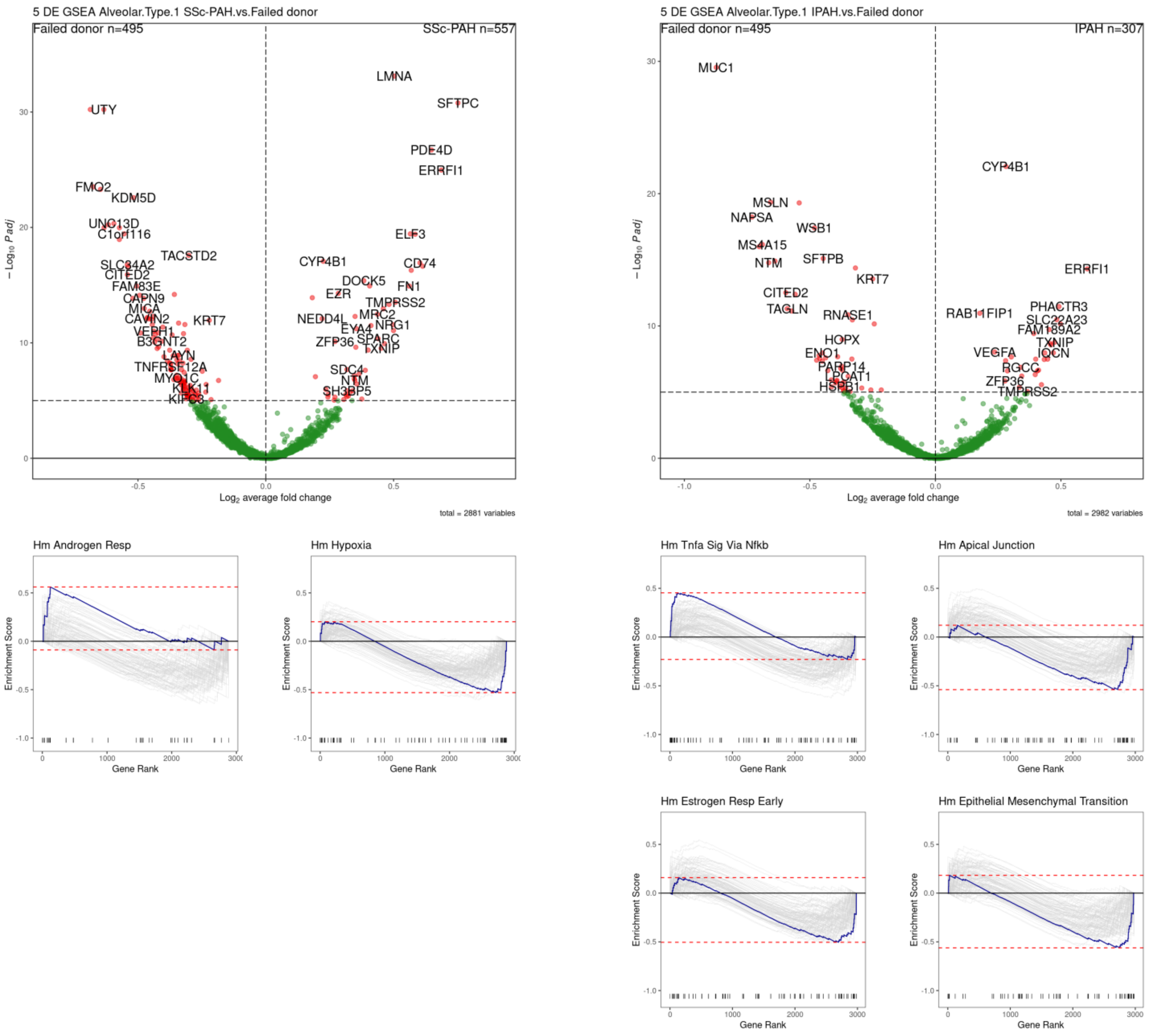


Supplemental Figure 10a

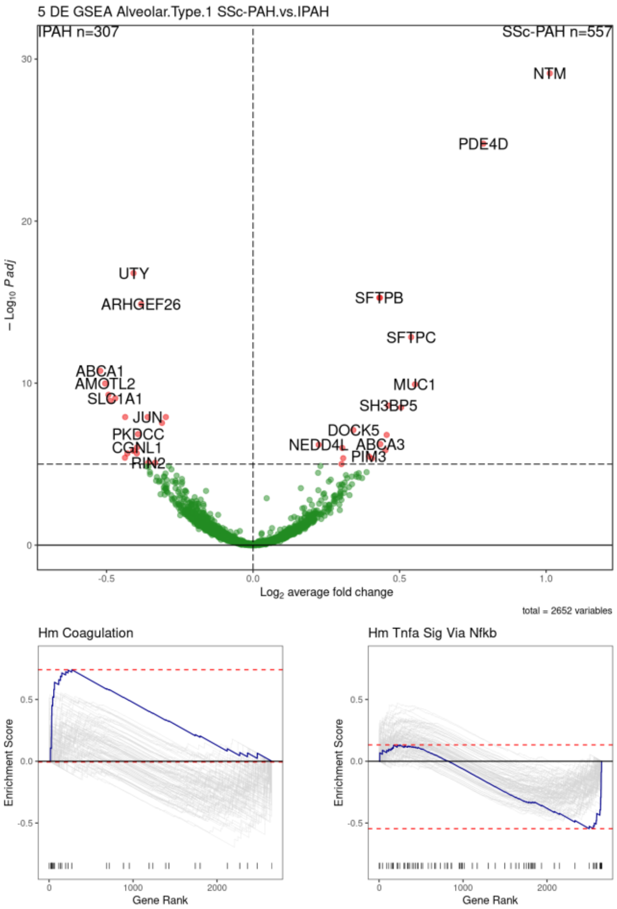


Supplemental Figure 10b

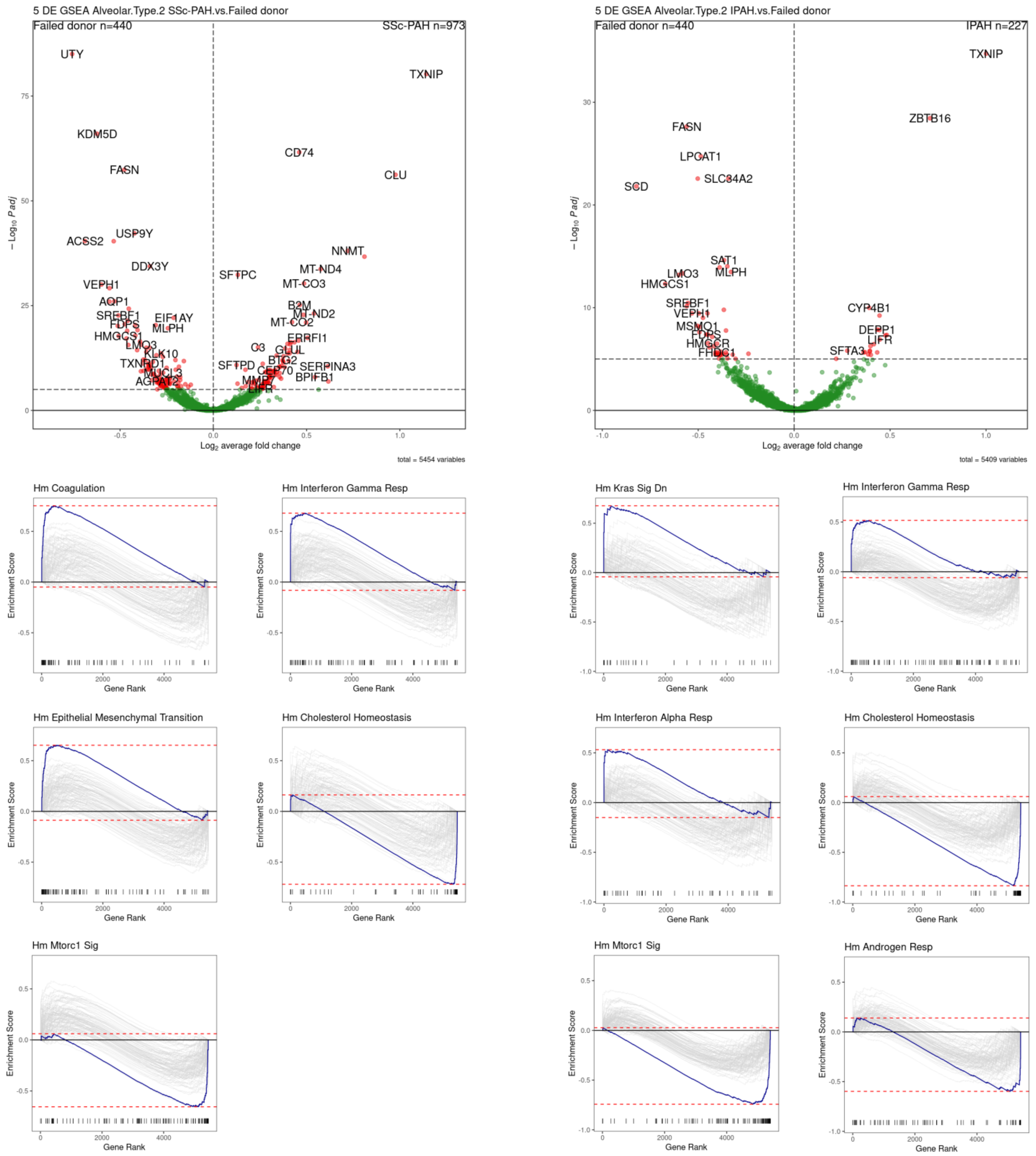


Supplemental Figure 11a

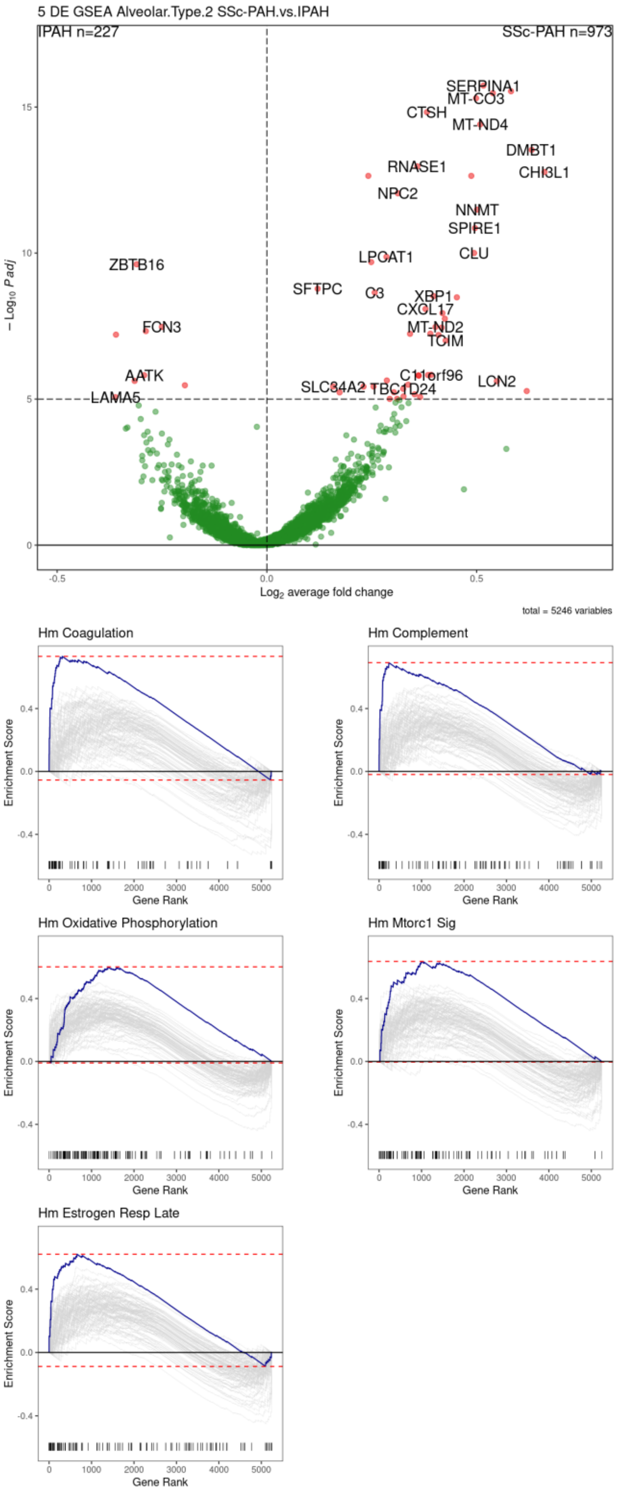


Supplemental Figure 11b

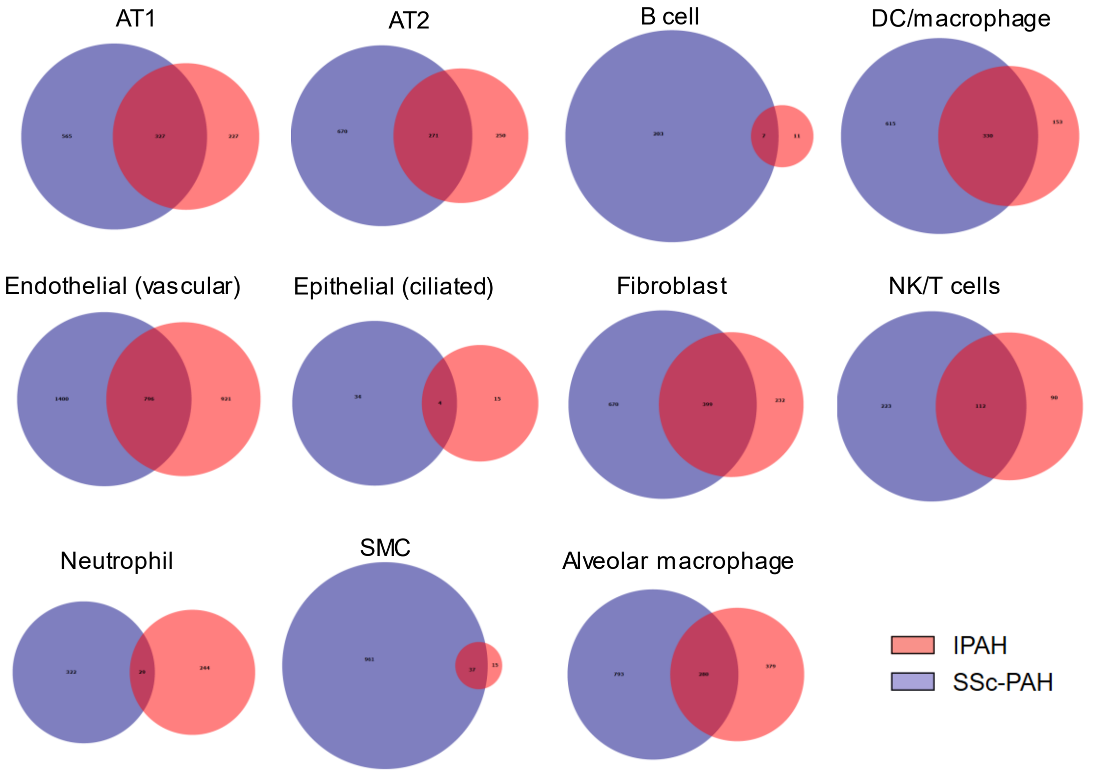


Supplemental Figure 12

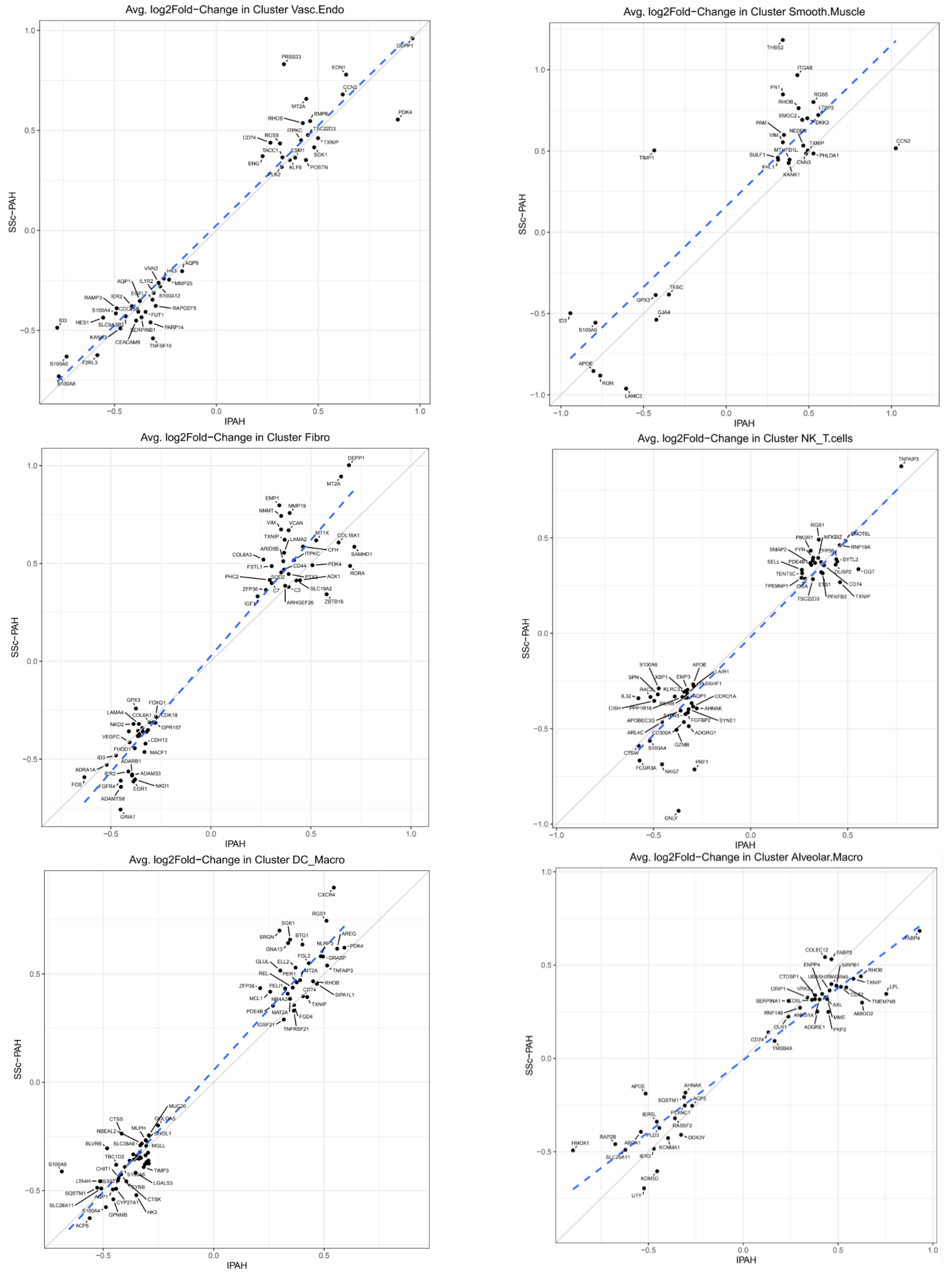


Supplemental Figure 13a

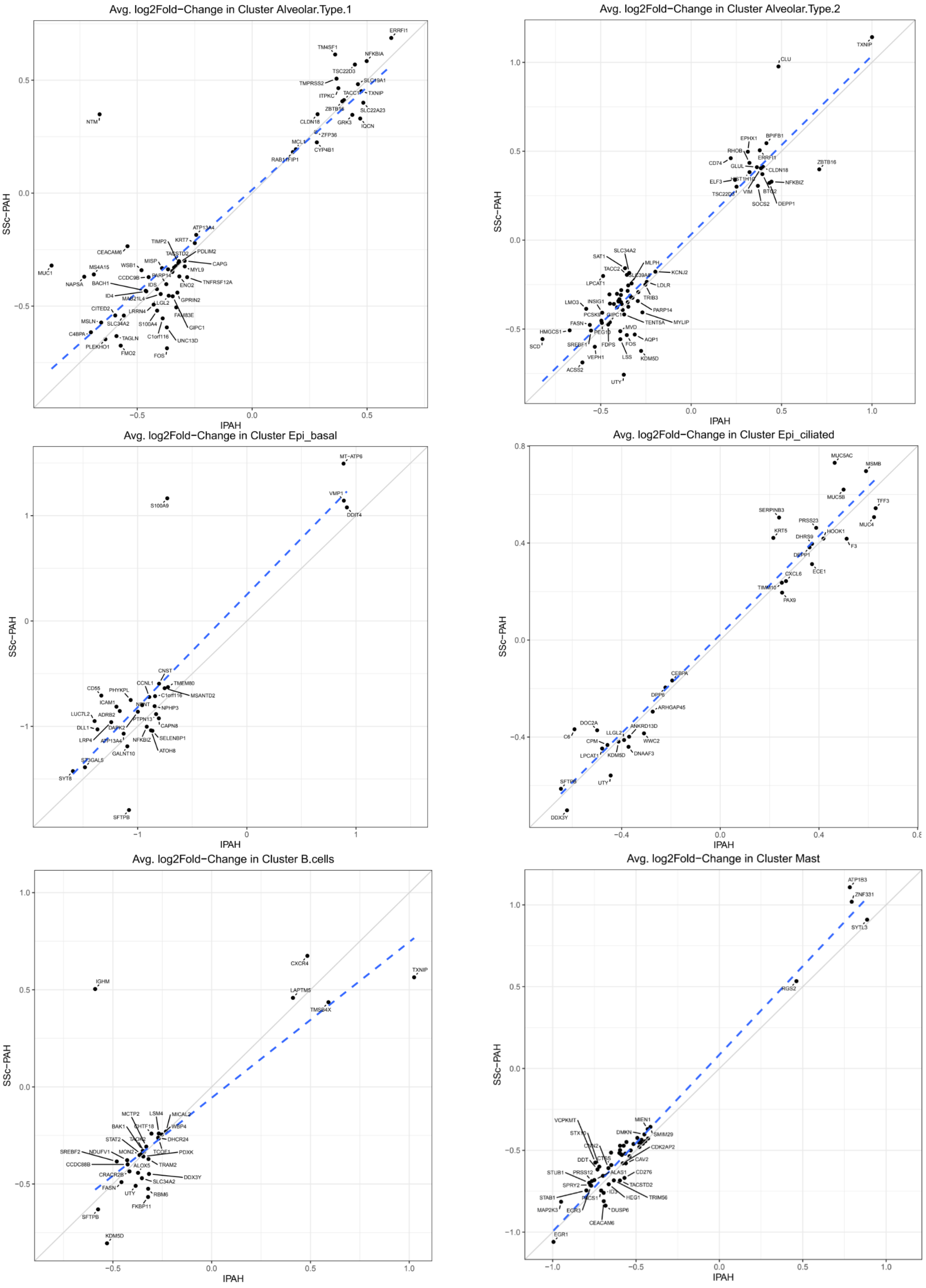


Supplemental Figure 13b

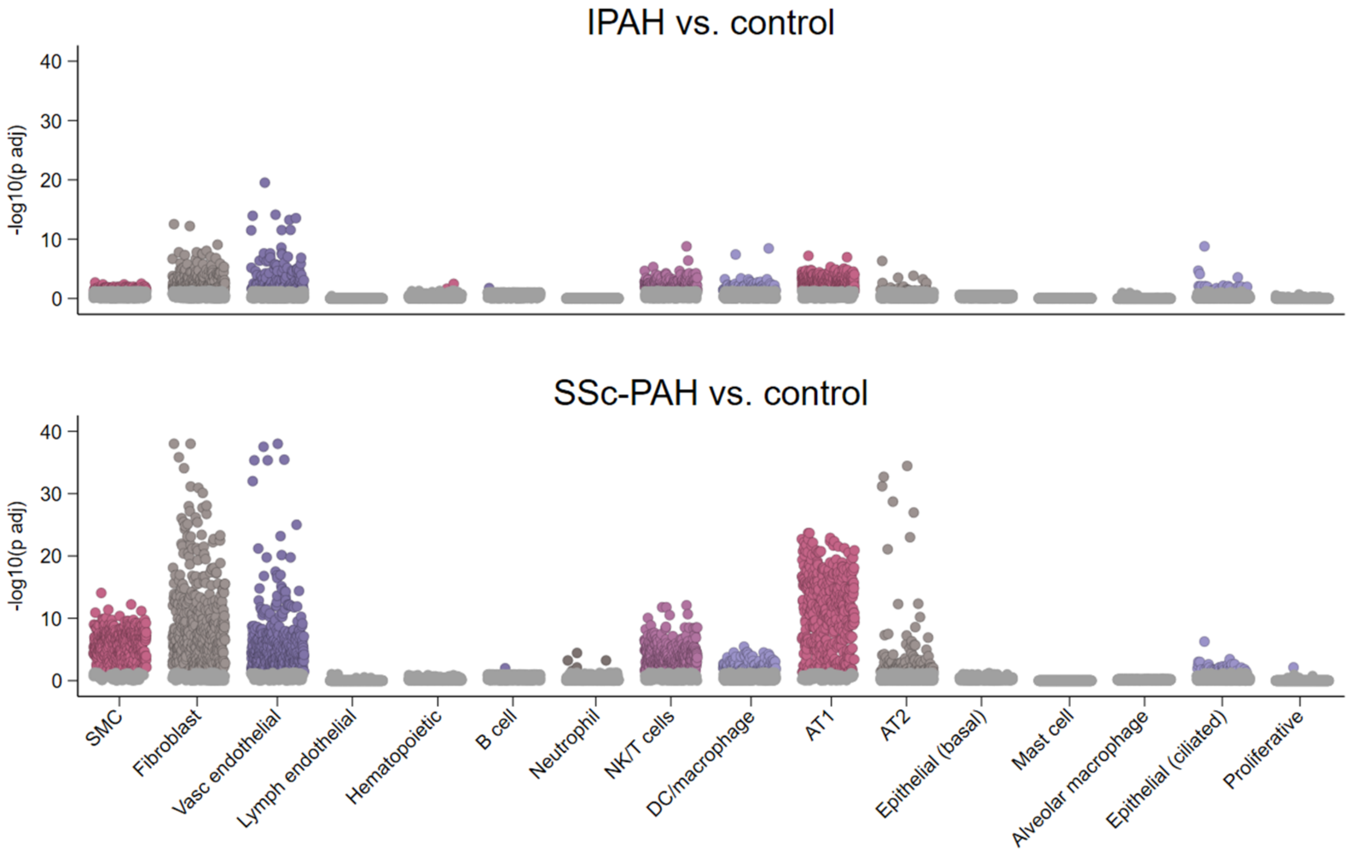


Supplemental Figure 14

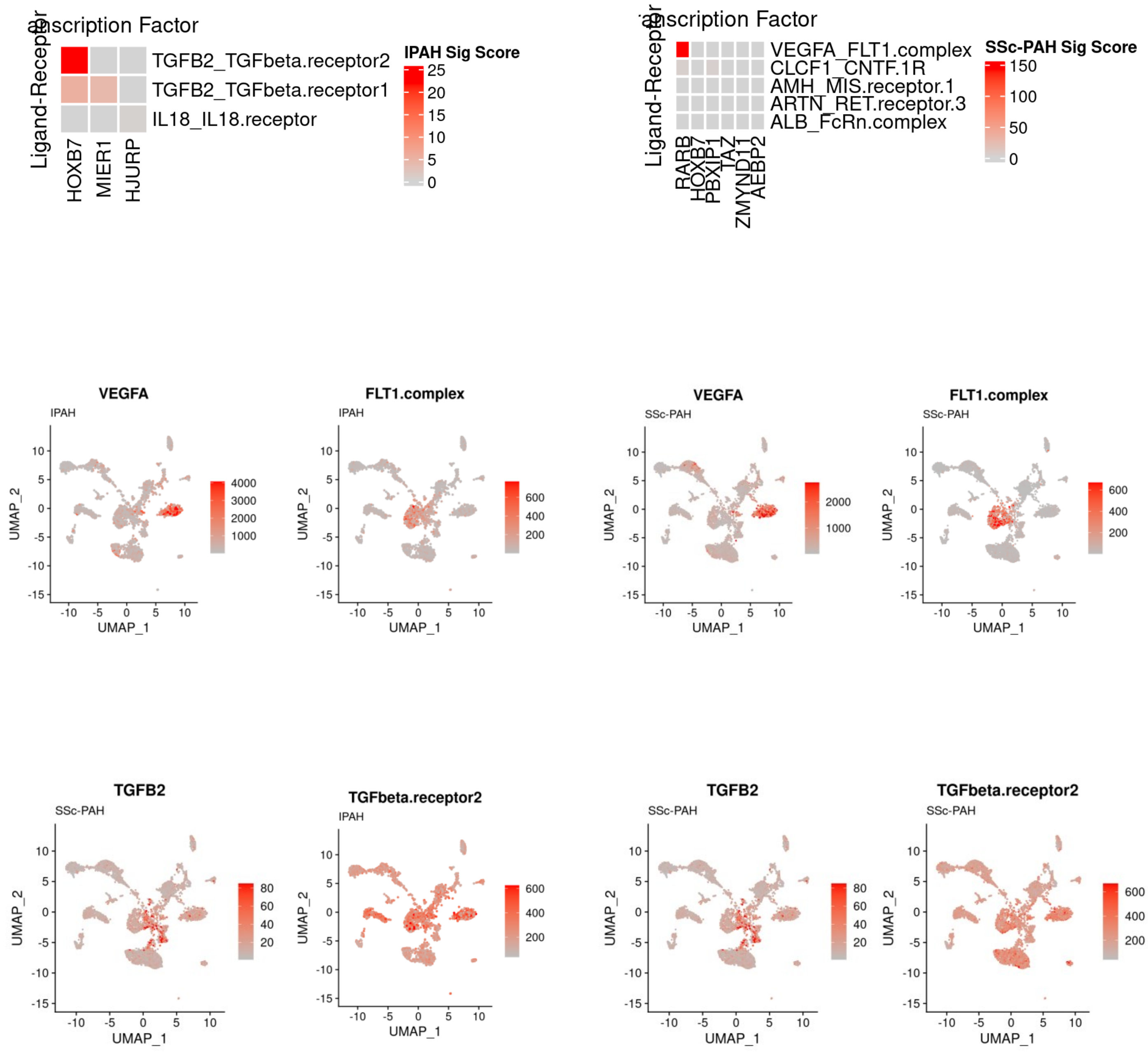


Supplemental Figure 15

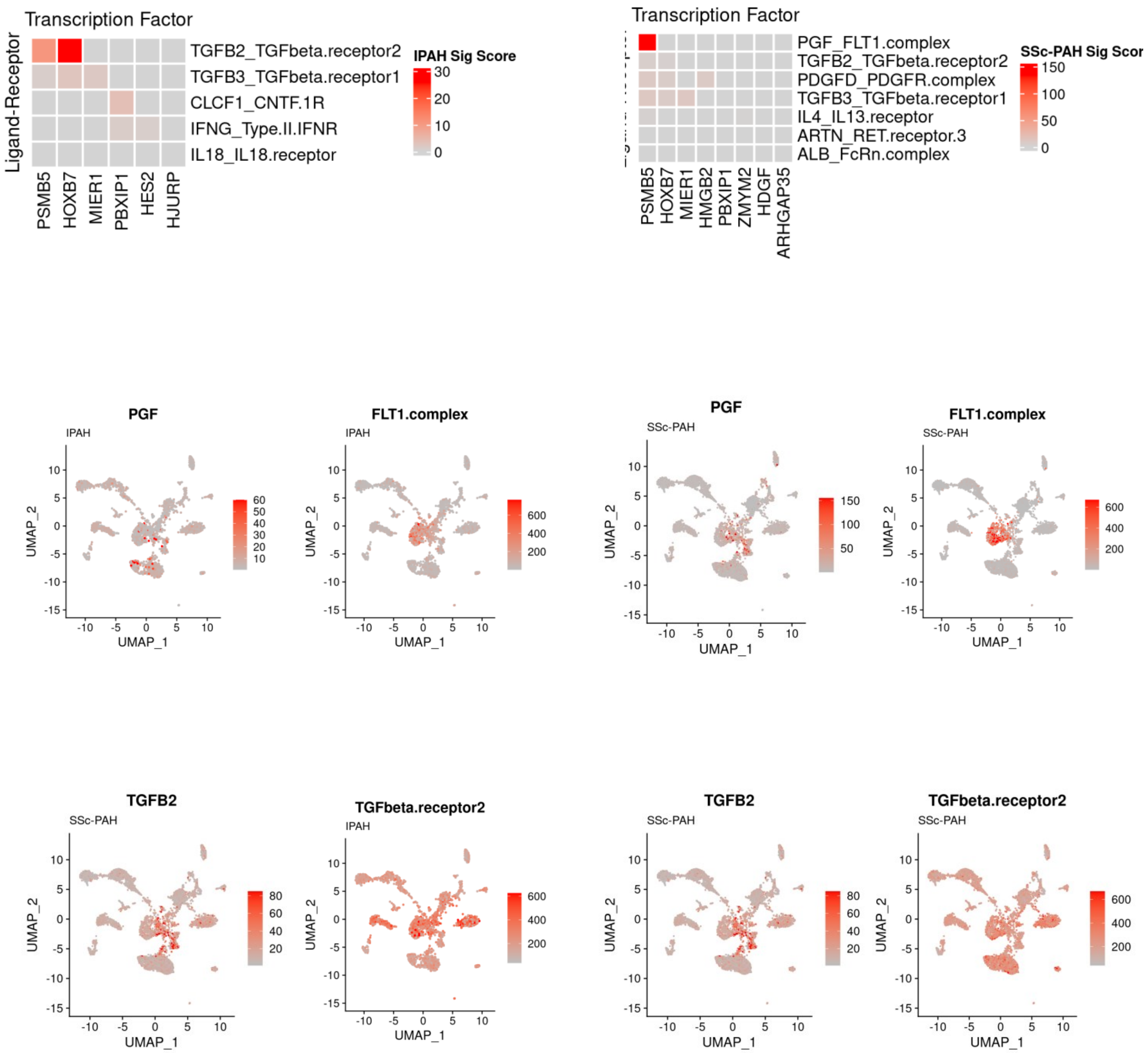


Supplemental Figure 16

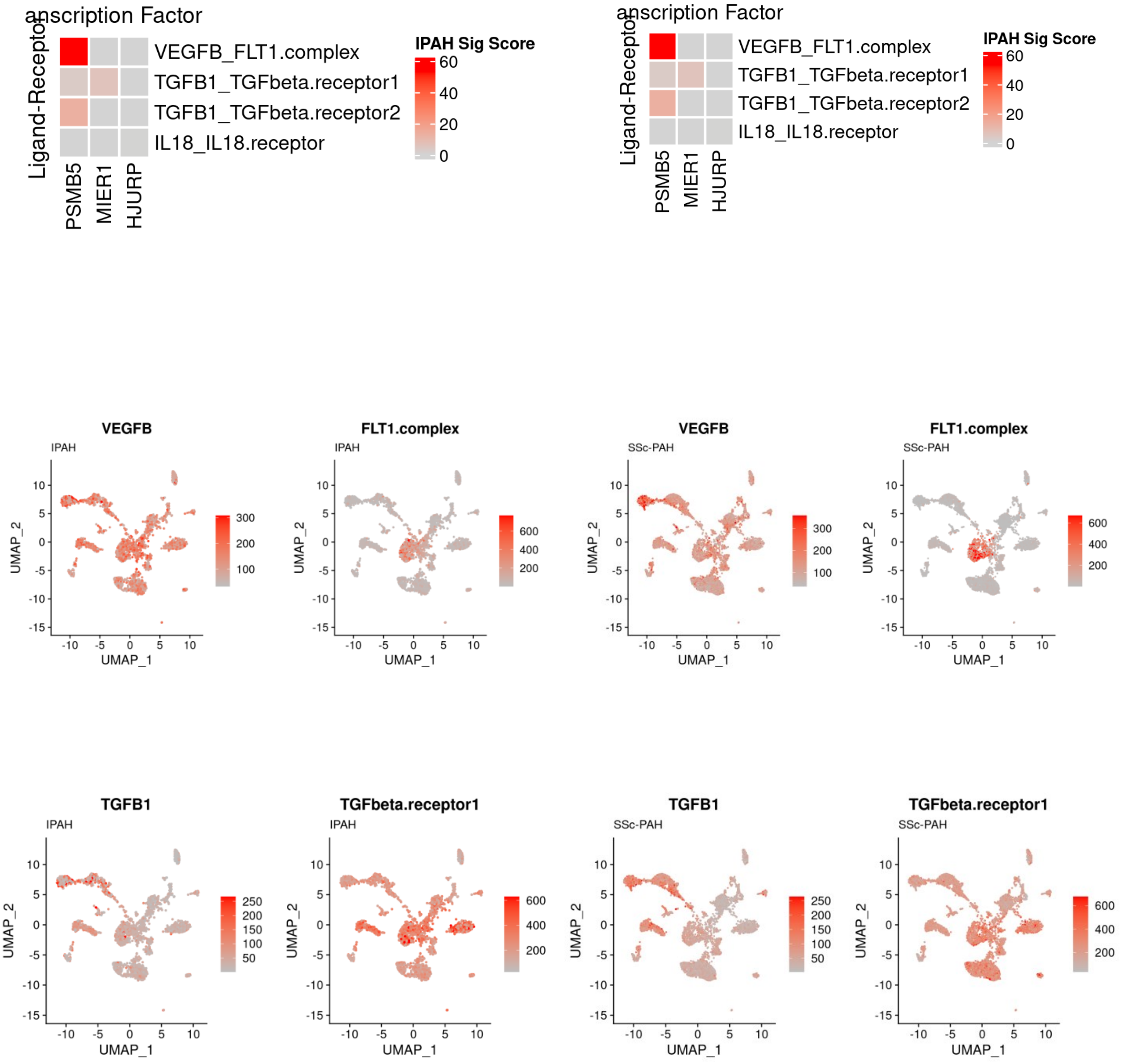


Supplemental Figure 17

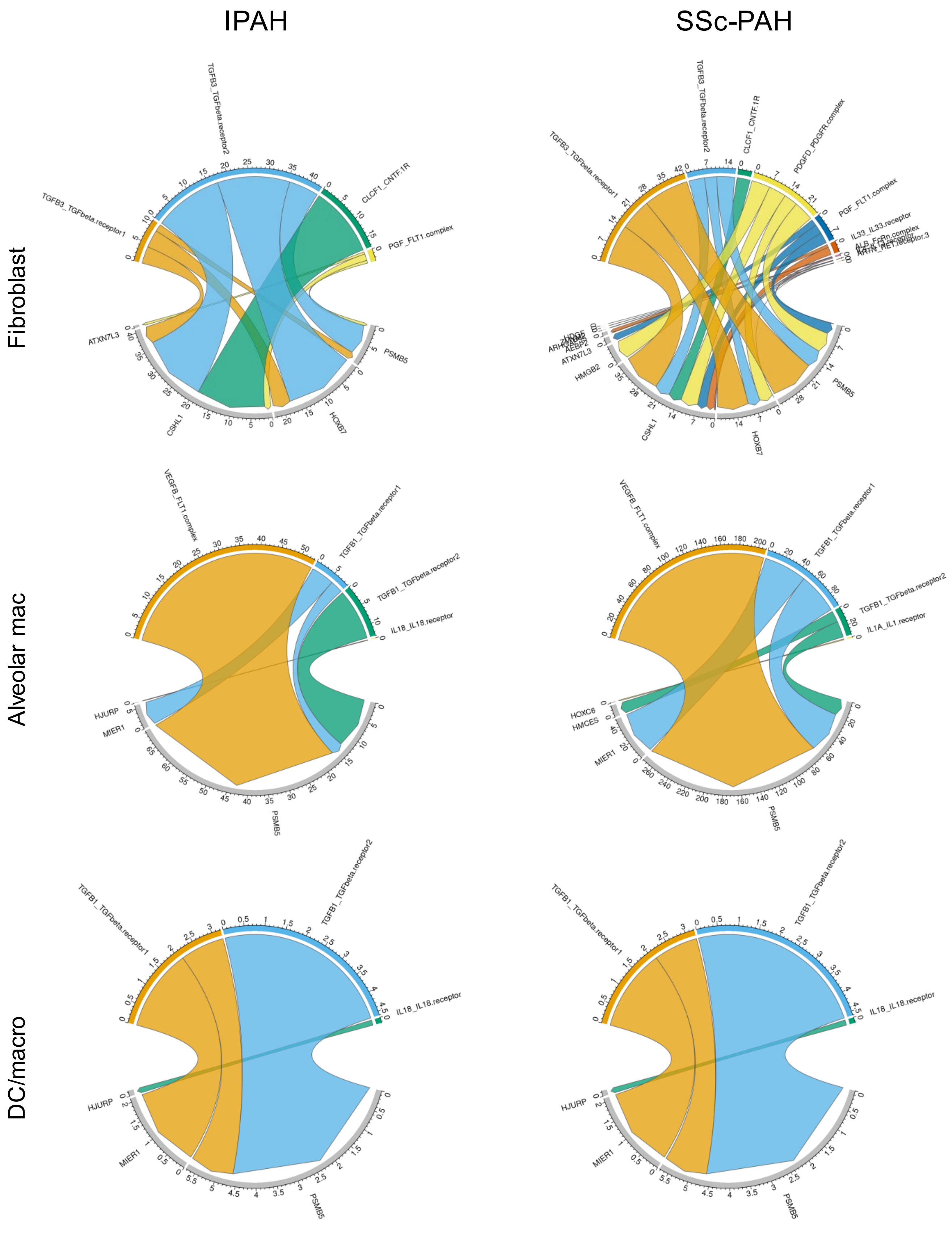


Supplemental Figure 18

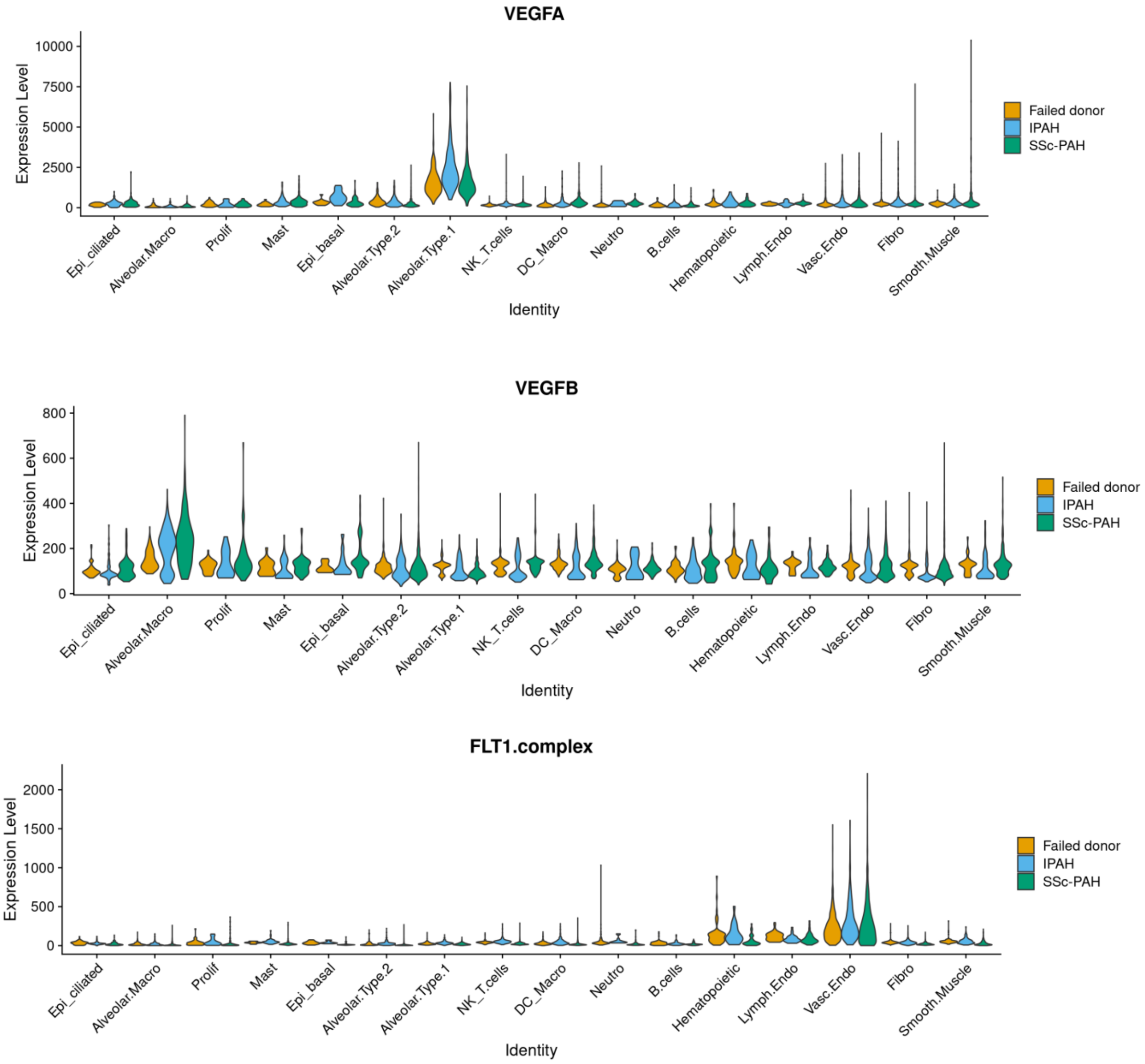


Supplemental Figure 19A

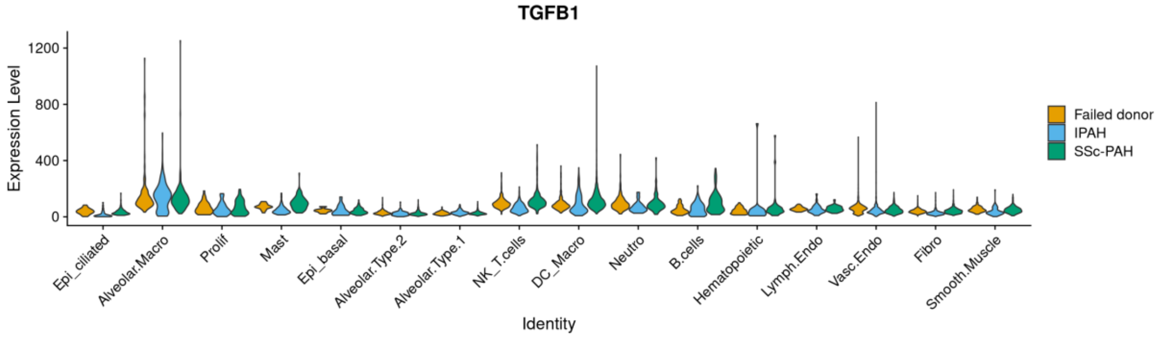


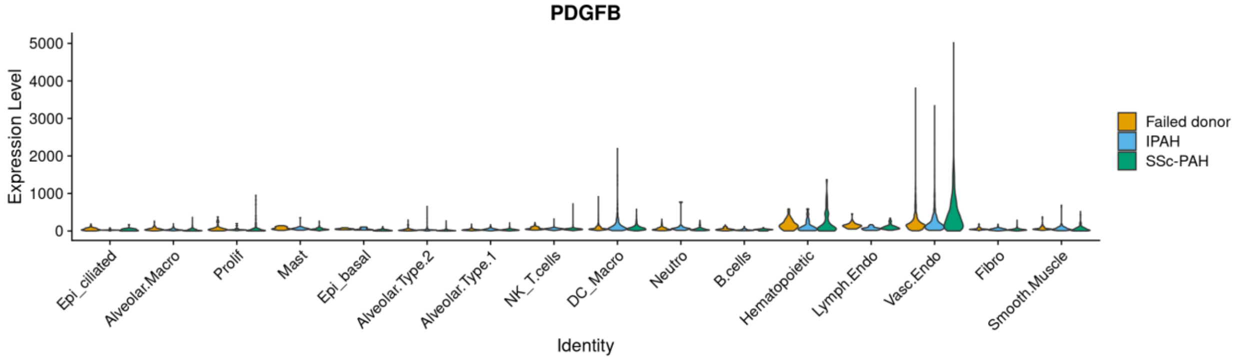


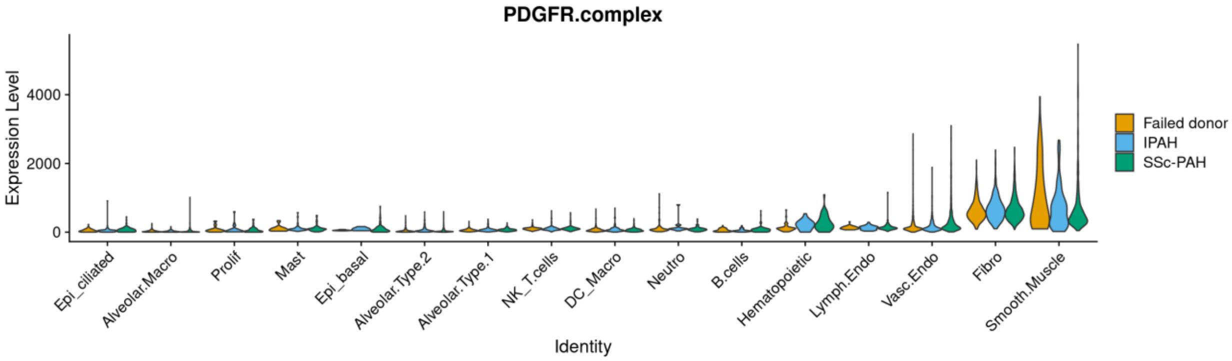


Supplemental Figure 19B

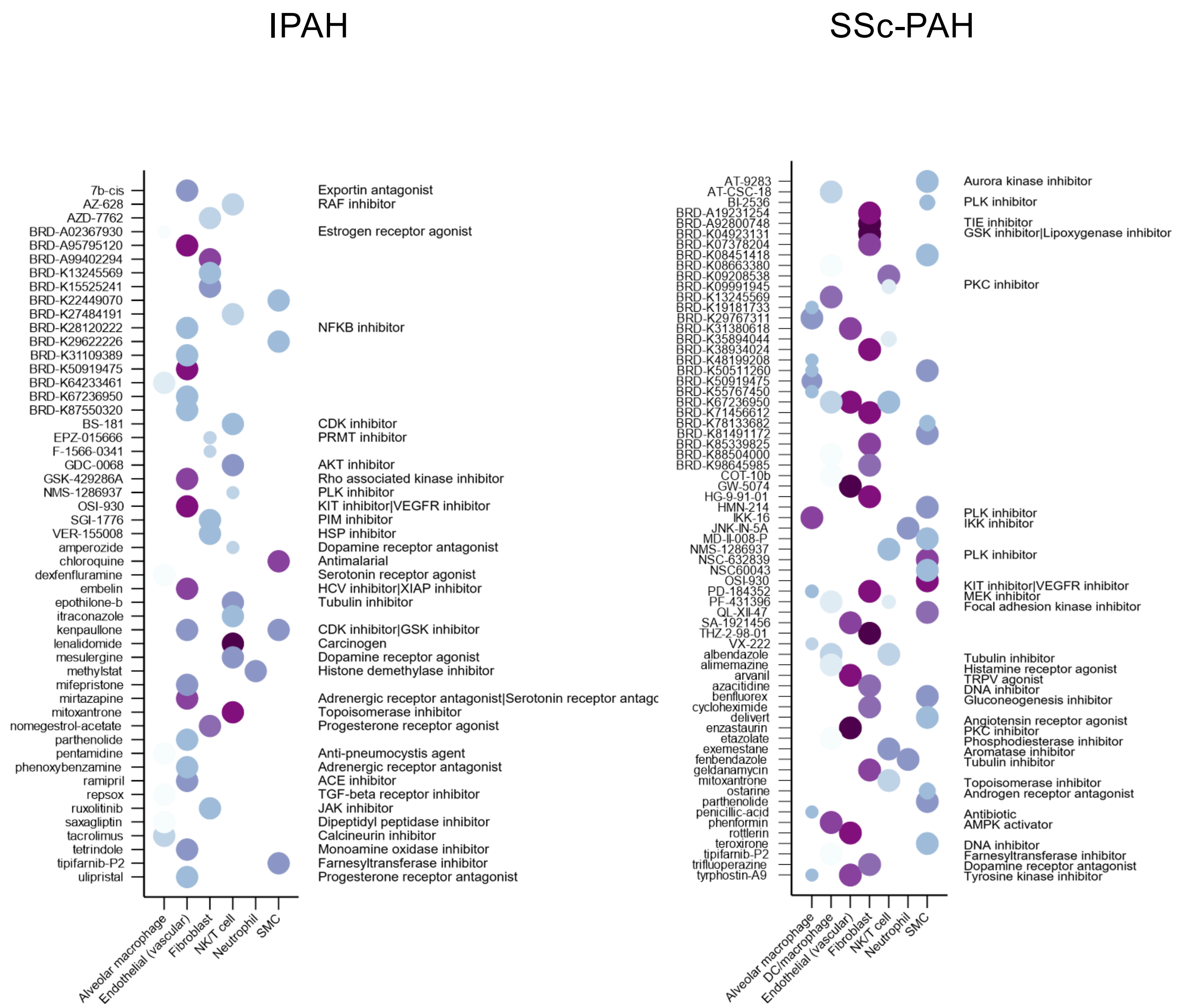


Supplemental Figure 20A

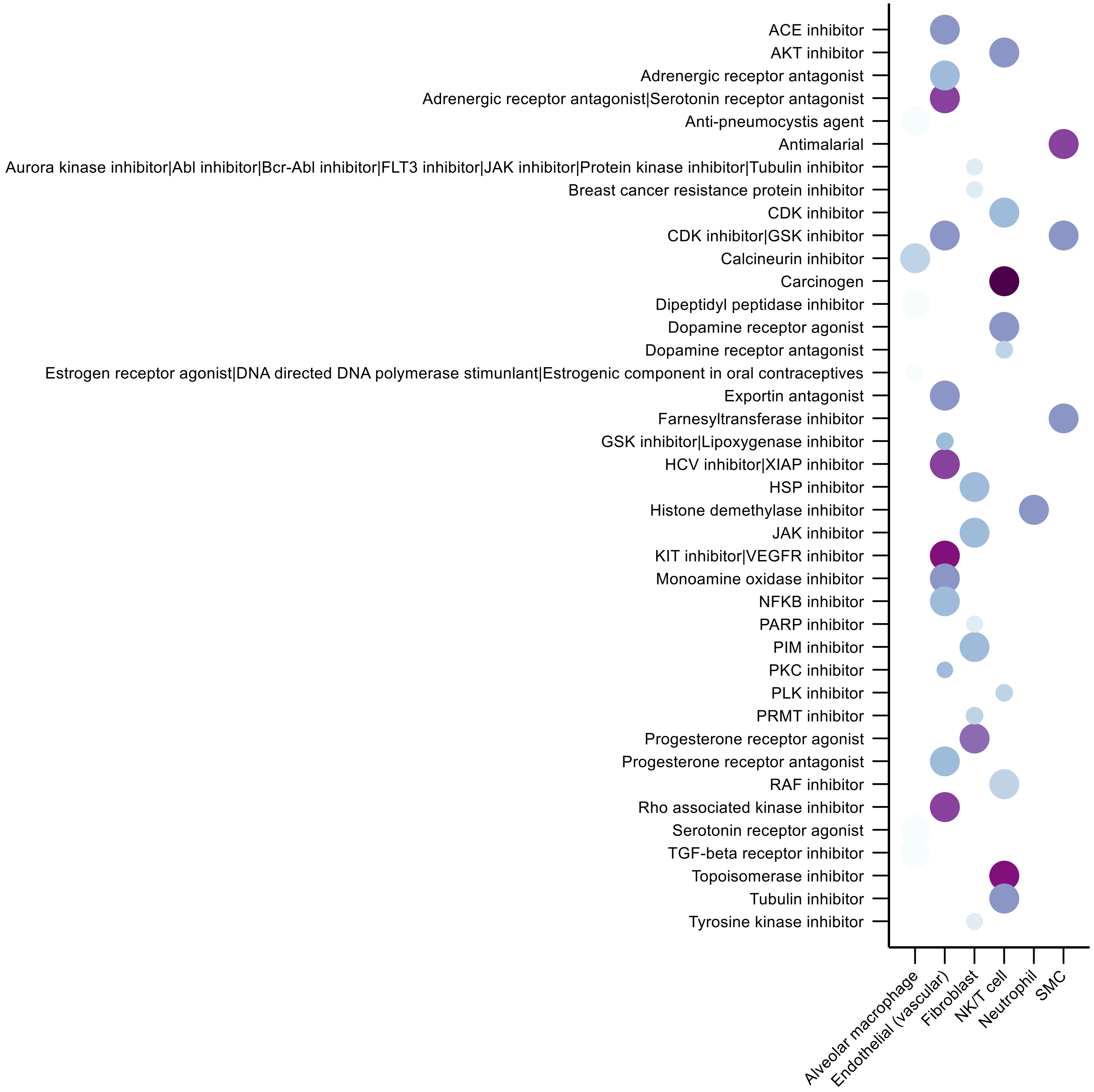


Supplemental Figure 20B

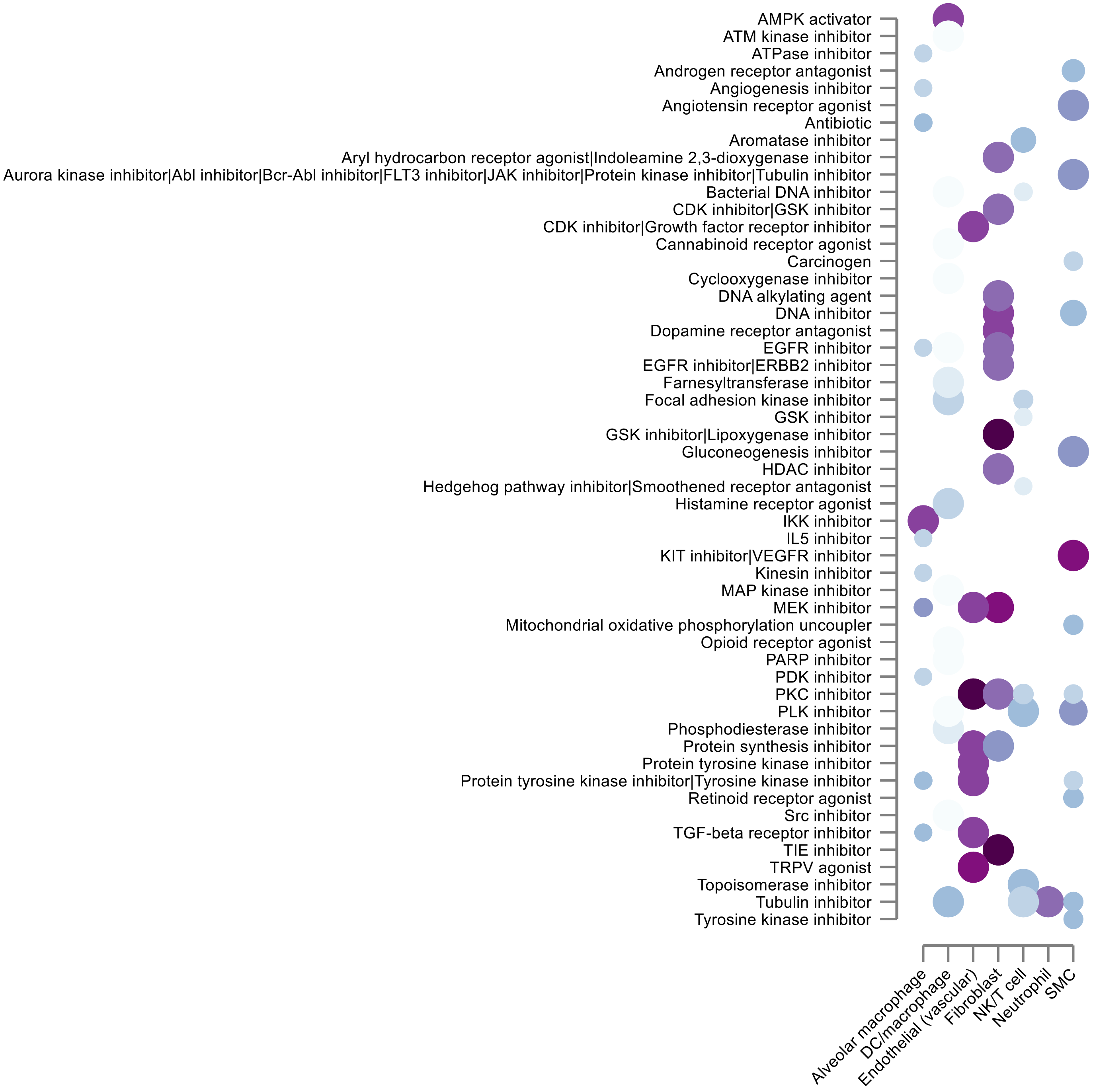


Supplemental Figure 20C
